# Midbrain dopaminergic inputs gate amygdala intercalated cell clusters by distinct and cooperative mechanisms

**DOI:** 10.1101/2020.10.29.360750

**Authors:** Ayla Aksoy-Aksel, Andrea Gall, Anna Seewald, Francesco Ferraguti, Ingrid Ehrlich

## Abstract

Dopaminergic signaling plays an important role in associative learning including fear and extinction learning. Dopaminergic midbrain neurons encode prediction error-like signals when threats differ from expectations. Within the amygdala, GABAergic intercalated cell (ITC) clusters receive the densest dopaminergic projections, but their physiological consequences are incompletely understood. ITCs are important for fear extinction, a function thought to be supported by activation of ventromedial cluster ITCs that inhibit central amygdala fear output. In mice, we reveal two distinct mechanisms how mesencephalic dopaminergic afferents control ITCs. Firstly, they co-release GABA to mediate rapid, direct inhibition. Secondly, dopamine suppresses inhibitory interactions between distinct ITC clusters via presynaptic D1-receptors. Early extinction training augments both, GABA co-release onto dorso-medial ITCs and dopamine-mediated suppression of dorso- to ventromedial inhibition between ITC clusters. These findings provide novel insights into dopaminergic mechanisms shaping the activity balance between distinct ITC clusters that could support their opposing roles in fear behavior.

## Introduction

Mesencephalic dopaminergic neurons in the ventral tegmental area (VTA) and substantia nigra pars compacta (SNC) constitute a neuromodulatory system that has been linked to error prediction and salience coding (Bromberg-Martin et al., 2010; Horvitz, 2000; Schultz et al., 1997). Prediction error signals drive the formation and the updating of stimulus-outcome associations by detection of mismatches between actual and expected experiences (Pearce and Hall, 1980; Rescorla and Wagner, 1972), which also operates when conditioned stimulus (CS)-unconditioned stimulus (US) and CS-no US associations need to be formed during fear conditioning and extinction learning, respectively (Bouton, 2004; McNally et al., 2011; Salinas-Hernandez et al., 2018). More specifically, omission of an expected aversive stimulus during the early sessions of extinction training activates a subset of VTA dopaminergic neurons (Salinas-Hernandez et al., 2018). Consequently, optogenetic inhibition of the VTA dopaminergic neurons during extinction training impairs, and their excitation enhances the extinction learning (Luo et al., 2018; Salinas-Hernandez et al., 2018).

The amygdala is a key structure that plays a critical role in mediating fear and anxiety related behaviors, and is a primary site for acquisition and storage of fear memory (Davis, 2000; Duvarci and Pare, 2014; Luo et al., 2018; Salinas-Hernandez et al., 2018). Dopamine (DA) is released in the amygdala during affective states such as stress or fear (Inglis and Moghaddam, 1999; Yokoyama et al., 2005) and direct pharmacological intervention on dopaminergic receptors in the central amygdala (CeA) or basal amygdala (BA) affects the acquisition and storage of fear memories (Guarraci et al., 1999; Lee et al., 2017).

Interestingly, selective optogenetic stimulation of midbrain dopaminergic axons has been shown to release not only the neuromodulator DA, but also fast-acting classical neurotransmitters such as GABA and/or glutamate in the dorsal striatum and nucleus accumbens (Granger et al., 2017; Tritsch et al., 2016). In the amygdala, glutamate was shown to be co-released with DA in CeA (Groessl et al., 2018; Mingote et al., 2015) and in BA, though preferentially onto fast-spiking interneurons (Lutas et al., 2019), a co-release of GABA has not been reported so far.

Several lines of evidence suggest that within the amygdala, the intercalated cells (ITCs) could be a key target for dopaminergic regulation. ITCs constitute a specialized network of GABAergic cells, and in mice are organized in several clusters around the basolateral complex of the amygdala (BLA) (Busti et al., 2011; Geracitano et al., 2007; Marowsky et al., 2005). Dopaminergic axons densely innervate the somata and dendrites of ITCs (Asan, 1997; Fuxe et al., 2003), however, the functional consequence of this innervation has been largely unexplored. Furthermore, ITCs show a high-expression level of pre- and post-synaptic DA-receptors (Fuxe et al., 2003; Pinto and Sesack, 2008; Wei et al., 2018). DA application in brain slices directly hyperpolarizes neurons in the medial and lateral ITC clusters via D1 receptors (DRD1) and thereby suppresses their output to CeA and BLA, respectively (Gregoriou et al., 2019; Manko et al., 2011; Marowsky et al., 2005).

In that respect, ITC clusters are ideally positioned to integrate dopaminergic signals with the sensory information that is either conveyed directly (Asede et al., 2015; Barsy et al., 2020; Strobel et al., 2015) or that has been preprocessed in the BLA (Herry and Johansen, 2014; Kwon et al., 2015; Pare et al., 2004). Indeed, ITCs and their plasticity have been shown to play a significant role in extinction (Amano et al., 2010; Likhtik et al., 2008) and more recently also in fear learning and memory (Asede et al., 2015; Busti et al., 2011; Huang et al., 2014; Kwon et al., 2015). The classical view on ITC function within amygdala circuits posits an inhibitory action onto neighboring CeA and BLA nuclei to gate information flow (Ehrlich et al., 2009; Marowsky et al., 2005; Morozov et al., 2011; Pare et al., 2004). However, there is emerging evidence pointing toward a more complex picture that involves temporal separation in the activity of the individual clusters, i.e. dorsomedial (dm)- and ventromedial (vm)-ITC clusters, in fear recall and extinction (Busti et al., 2011). Considering the connections between the different clusters (Asede et al., 2015; Busti et al., 2011; Duvarci and Pare, 2014; Royer et al., 2000), it is plausible that ITCs’ inhibitory actions not only on CeA and BLA, but also onto other ITC clusters could be modulated by DA.

Thus, while mounting evidence supports that midbrain dopaminergic inputs play important roles in controlling amygdala function in fear learning as well as extinction (Abraham et al., 2014; Lee et al., 2017), the cellular impact onto specific amygdala microcircuits is incompletely understood (Grace et al., 2007; Lee et al., 2017). To address this knowledge gap, we used specific optogenetic stimulation of dopaminergic axons from VTA/SNC to explore the functional impact onto ITCs. We observed co-release of GABA from dopaminergic axons onto ITCs mediating fast inhibition. Phasic stimulation of dopaminergic fibers suppressed spontaneous inhibitory inputs onto dm-ITCs. Consequently, when interrogating transmission between ITCs clusters, we observed a presynaptic depression mediated by DRD1. Extinction learning promoted the co-release of GABA in dm-ITCs and the DA-induced suppression of inhibition between dm- and vm-ITC clusters. Our results demonstrate a dual mechanism of action of dopaminergic inputs with a cooperative fast ionotropic and a slow metabotropic component, that jointly regulates the inhibitory network formed between ITCs. We suggest that this can tip the activity balance between individual clusters to support fear suppression.

## Results

### Dopaminergic fibers from VTA/SNC innervate ITC clusters in amygdala

We first examined the projections of midbrain dopaminergic inputs to amygdala ITC clusters and surrounding amygdala regions. To this end, we targeted dopaminergic cells selectively using a DAT-IRES-Cre mouse line and injected a Cre-dependent recombinant adenoassociated virus (AAV) encoding ChR2-YFP into the VTA/SNC (Figure 1–figure supplement 1A). Our injections resulted in expression of ChR2-YFP mostly in VTA and to a lesser extent in SNC neurons, the vast majority of which were immunopositive for tyrosine hydroxylase (TH) (Figure 1–figure supplement 1B-C). We next examined YFP+ fibers in the amygdala (Figure 1–figure supplement 2A). YFP+ fibers targeting FoxP2-positive neurons in the dm- and vm-ITC clusters were also immunoreactive for TH, indicating that they are dopaminergic in nature (Figure 1A-B). In line with previous reports (Lutas et al., 2019; Mingote et al., 2015), we also observed YFP+ fibers colocalized with TH-labeled axons in the CeA and more sparsely in the BA (Figure1–figure supplement 2B-C).

**Figure 1.**
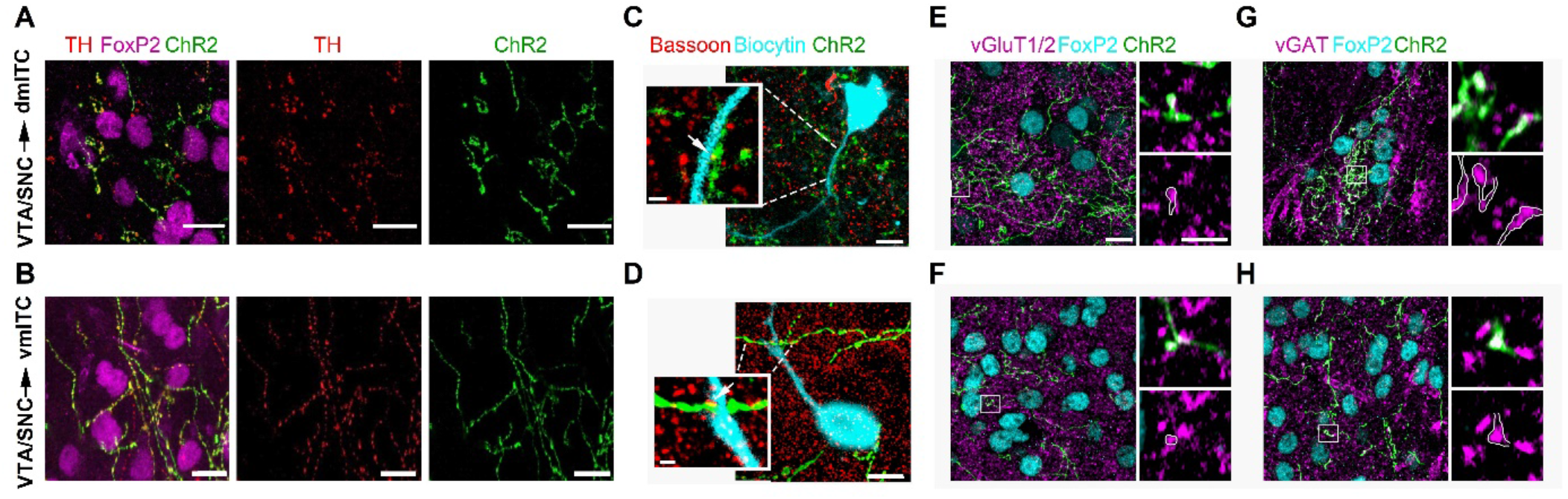
Dopaminergic fibers from VTA/SNC targeting dm- and vm-ITC clusters in the amygdala co-label with vGAT and vGluT1/2. **(A-B)** Maximum intensity projection confocal images illustrating ChR2-YPF axons (green) originating from TH-positive dopaminergic neurons (TH, red) in the VTA/SNC targeting FoxP2-positive neurons (magenta) in the dm- and vm-ITC clusters. Overlay (left), TH staining (middle), ChR2-YPF (left). **(C-D)** Overlay confocal images of ChR2-YFP labeled axons (green) and biocytin filled cells (turquoise) in dm-ITC and vm-ITC clusters immunostained for the presynaptic marker Bassoon to examine putative active zones. **(E-F)** Confocal images of dm-ITC and vm-ITC clusters including FoxP2 positive neurons (turquoise) that contain ChR2-YFP fibers (green) co-labeled for the presynaptic markers vGluT1/vGluT2 (magenta). Right panels show a higher magnification of the boxed area containing an example of a bouton co-expressing ChR2-YFP and vGluT1/vGluT2 outlined in white. **(G-H)** Confocal images of dm-ITC and vm-ITC clusters including FoxP2 positive neurons (turquoise) that contain ChR2-YFP fibers (green) co-labeled for the presynaptic marker vGAT (magenta). Right panels show a higher magnification of the boxed area containing an example of a bouton co-expressing ChR2-YFP and vGAT outlined in white. Thickness of confocal z-stacks: (A) 11.1 μM; (B) 12.5 μM; (C) 8.06 μM and 2.01 μM for the cell and synapse on dendrite, respectively; (D) 12.2 μM and 2.2 μM for the cell and synapse on dendrite, respectively; (E-H) left panels 8.83 μm, right panels single plane of 0.18 μm nominal thickness. Scale bars: (A-B) 10 μm, (C-D) 5 μm and 1 μM, (E-H) 10 μm for left panels, 3 μm for right panels.

We next asked if YFP+ midbrain afferents made synaptic contacts with neurons in the dm- and vm-ITC clusters. To this aim, we filled ITCs with biocytin, which allowed us to visualize their somato-dendritic domain. We then labeled putative presynaptic active zones by immunostaining for Bassoon (Liu et al., 2018). Colocalization of Bassoon and ChR2-YFP was observed in close proximity to dendrites and somata of ITCs in dm- and vm-clusters (Figure 1C-D), suggesting that DA midbrain inputs make functional synaptic contacts.

Midbrain dopaminergic neurons from VTA and SNC co-release GABA and/or glutamate in the dorsal striatum and nucleus accumbens (Granger et al., 2017; Mingote et al., 2015; Stuber et al., 2010; Tecuapetla et al., 2010; Tritsch et al., 2012; Tritsch et al., 2016). In the amygdala, functional co-release of glutamate from dopaminergic VTA afferents has been demonstrated in CeA (Mingote et al., 2015) and to a lesser extent in the BA (Lutas et al., 2019). To adress whether this may also be the case for ITCs, we first checked for the presence of vesicular GABA or glutamate transporters in ChR2-YFP+ axons. We indeed observed immunoreactivity for the vesicular glutamate transporters vGluT1/2 and vesicular GABA transporter vGAT in a subset of ChR2-YFP+ presynaptic boutons within FoxP2-positive dm- and vm-ITC clusters (Figure 1E-H), corroborating the hypothesis of GABA and glutamate co-release from midbrain dopaminergic fibers. Because of incomplete penetration of the vGluT1/2 and vGAT antibodies into slices, a quantitative analysis of the colocalization with ChR2-YFP+ terminals was not carried out.

### Dopaminergic fibers co-release mainly GABA in the medial ITC clusters

In light of our anatomical observations of a possible co-release of GABA or glutamate, we explored whether optical stimulation of ChR2-YFP+ fibers from confirmed dopaminergic midbrain injections (Figure2–figure supplement 1A-C) evoked fast postsynaptic currents (PSCs) in dm- and vm-ITCs. Indeed, we could detect PSCs in approximately 50-60% of the neurons recorded in whole cell patch clamp mode in ITC clusters as well as in the CeA (Figure 2A-B, D, G), but not in principal neurons of the BA (10 neurons recorded from 5 animals showing co-release in either CeA or ITCs, Figure 2–figure supplement 2D).

**Figure 2.**
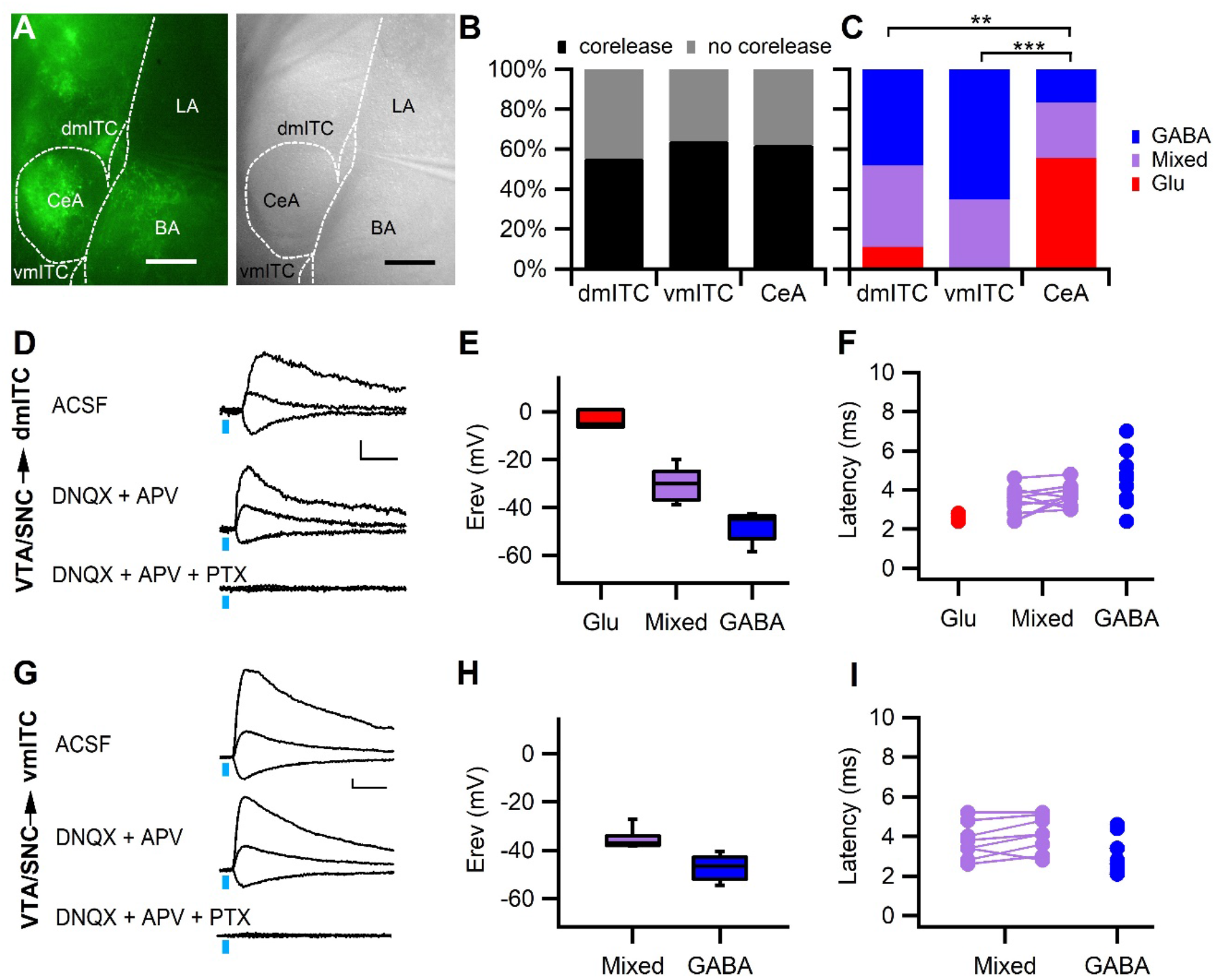
Stimulation of midbrain dopaminergic fibers evokes mainly GABAergic PSCs in dm- and vm-ITCs. **(A)** Fluorescence (left) and infrared-DIC image of an amygdala slice with patch recording pipette in the dm-ITC cluster (right). ChR2-YFP expressing axons from dopaminergic midbrain were observed in CeA, amygdalostriatal transition zone, BA, and ITCs. **(B)** Fast postsynaptic currents (PSCs) were evoked by stimulation of dopaminergic fibers in a comparable fraction of recorded ITCs and CeA neurons (Fisher’s exact test = 0.763, p=0.691; 55% for dm-ITCs, n=49; 64% for vm-ITC, n=36; and 62% for CeA, n=29). **(C)** Distribution of fast PSCs in dm-ITCs, vm-ITCs, and CeA neurons by response type. ITCs show mostly GABAergic PSCs (dm-ITCs, n=27: GABA 48%, mixed 41%, Glu 11%; vm-ITCs, n=23: GABA 65%, mixed 35%,), whereas CeA neurons (n= 18) exhibited mostly glutamatergic PSCs (GABA 17%, mixed 28%, Glu 55%). Response types were significantly different between all regions (Fisher’s exact test = 21.41, p<0.001). A pairwise comparison revealed differences between CeA and ITC clusters (CeA vs. dm-ITC **p=0.005; for CeA vs. vm-ITC ***p<0.001). **(D and G)** Representative traces of light-evoked mixed PSCs recorded at −70 mV, 0 mV, and 40 mV from a dm-ITC (D) and a vm-ITC (G). Application of glutamate-receptor blockers DNQX (20 μM) and APV (100 μM) had a small effect, whereas the GABA_A_-R channel blocker PTX (100 μM) abolished the PSCs entirely. Scale bars: 10 pA, 10 ms. **(E and H)** Box plot of PSC reversal potentials (Erev) for dm-ITCs (E) and vm-ITCs (H) by PSC type. **(E)** Erev in dm-ITCs was −3.49 ± 2.21 mV for glutamatergic PSCs, −48.15 ± 1.65 mV for GABAergic PSCs, and −30.21 ± 2.07 mV for mixed PSCs (one-way ANOVA, F (2, 24) = 70.563, p<0.001). **(H)** Erev in vm-ITCs was −47.12 ± 1.31 mV for GABAergic PSCs and −35.57 ± 1.32 mV for mixed PSCs (unpaired t-test p<0.001). **(F and I)** Latencies of PSCs were consistent with monosynaptic connections. **(F)** Latencies in dm-ITCs were 2.60 ± 0.12 ms (n=3) for pure glutamatergic, 3.45 ± 0.21 ms for the glutamatergic and 3.67 ± 0.16 ms (n=11) for the GABAergic components of mixed PSCs, and 4.33 ± 0.35 ms (n=13) for pure GABAergic PSCs. **(I)** Latencies in vm-ITCs were 3.75 ± 0.32 ms for glutamatergic and 4.09 ± 0.32 ms (n=8) for GABAergic components of mixed PSCs, and 3.04 ± 0.25 ms (n=15) for pure GABAergic PSCs.

To classify fast PSCs according to neurotransmitter type, we first determined their reversal potential (Figure 2E and H). In a large fraction of neurons within dm- and vm-ITC clusters, we found PSCs that reversed close to the calculated reversal potential for chloride (−47.3 mV), suggesting that they were GABAergic (Figure 2C, E and H).

Another fraction of neurons displayed PSCs with more depolarized reversal potentials (>-40 mV and <-15mV), suggesting that they were mixed PSCs, and finally a small fraction of dm-ITCs displayed PSCs that reversed close to 0mV, suggesting that they were glutamatergic (Figure 2C, E and H). To ascertain our classification, we identified the neurotransmitter receptors involved using pharmacological blockers of GABA_A_-receptors, as well as AMPA and NMDA-type glutamate receptors in a subset of the recorded ITC neurons (Figure 2D, G). Applying the same approach to CeA neurons, in accordance with previous reports, we detected glutamatergic PSCs (Mingote et al., 2015), but also GABAergic and mixed PSCs (Figure 2C and Figure 2–figure supplement 2A-B). Overall, the relative contribution of neurotransmitters mediating the fast PSCs significantly differed between neurons in dm-ITC or vm-ITC clusters compared with CeA neurons (Figure 2C), with the latter having a higher proportion of glutamatergic PSCs.

In line with a monosynaptic innervation, all components of the recorded inhibitory and excitatory PSCs had short latencies in both ITC clusters and CeA (Figure 2F and I, Figure 2–figure supplement 2C). In dm-ITCs, we also directly confirmed that PSCs were monosynaptic by demonstrating that they were abolished by the sodium channel blocker tetrodotoxin (TTX) and recovered in the presence of TTX and the potassium channel blocker 4-aminopyridine (4-AP) (Figure 2–figure supplement 3A-B) (Petreanu et al., 2009). Taken together, our results strongly suggest that projections originating from VTA and SNC dopaminergic neurons mainly co-release GABA in medial ITC clusters to induce GABA_A_-receptor mediated fast inhibitory PSCs (IPSCs).

### Phasic stimulation of dopaminergic fibers alters sIPSC amplitude in dm-ITCs

Phasic activity of midbrain dopaminergic neurons has been shown to release dopamine in dorsal striatum and nucleus accumbens both *ex vivo* and *in vivo* (Schultz et al., 1997; Stuber et al., 2010). If this is also the case for ITCs, we should be able to detect an electrophysiological signature or a modulatory effect of dopaminergic inputs onto ITCs. We addressed this by stimulating dopaminergic midbrain afferents with 30 Hz light pulse trains to simulate phasic activity, while comparing the frequency and amplitude of spontaneous synaptic inputs (sPSCs) onto dm-ITCs before and after stimulation (Figure 3A). While we did not detect changes in the frequency for either sIPSCs or sEPSCs (Figure 3A-B), we observed a small but significant reduction in the amplitude of sIPSCs, but not sEPSCs (Figure 3C-D). Conversely, tonic 1 Hz stimulation had no effect on frequency or amplitude of sIPSCs (Figure 3–figure supplement 1A-B). Stimulation of dopaminergic midbrain inputs at 30 Hz also decreased the amplitude of sIPSC in vm-ITCs (Figure3–figure supplement 2A-B), corroborating our findings from dm-ITCs. Although we cannot completely rule out contributions from co-released transmitters, our results strongly suggest that phasic release of DA by midbrain neurons has a neuromodulatory effect on ITC inhibitory inputs.

**Figure 3.**
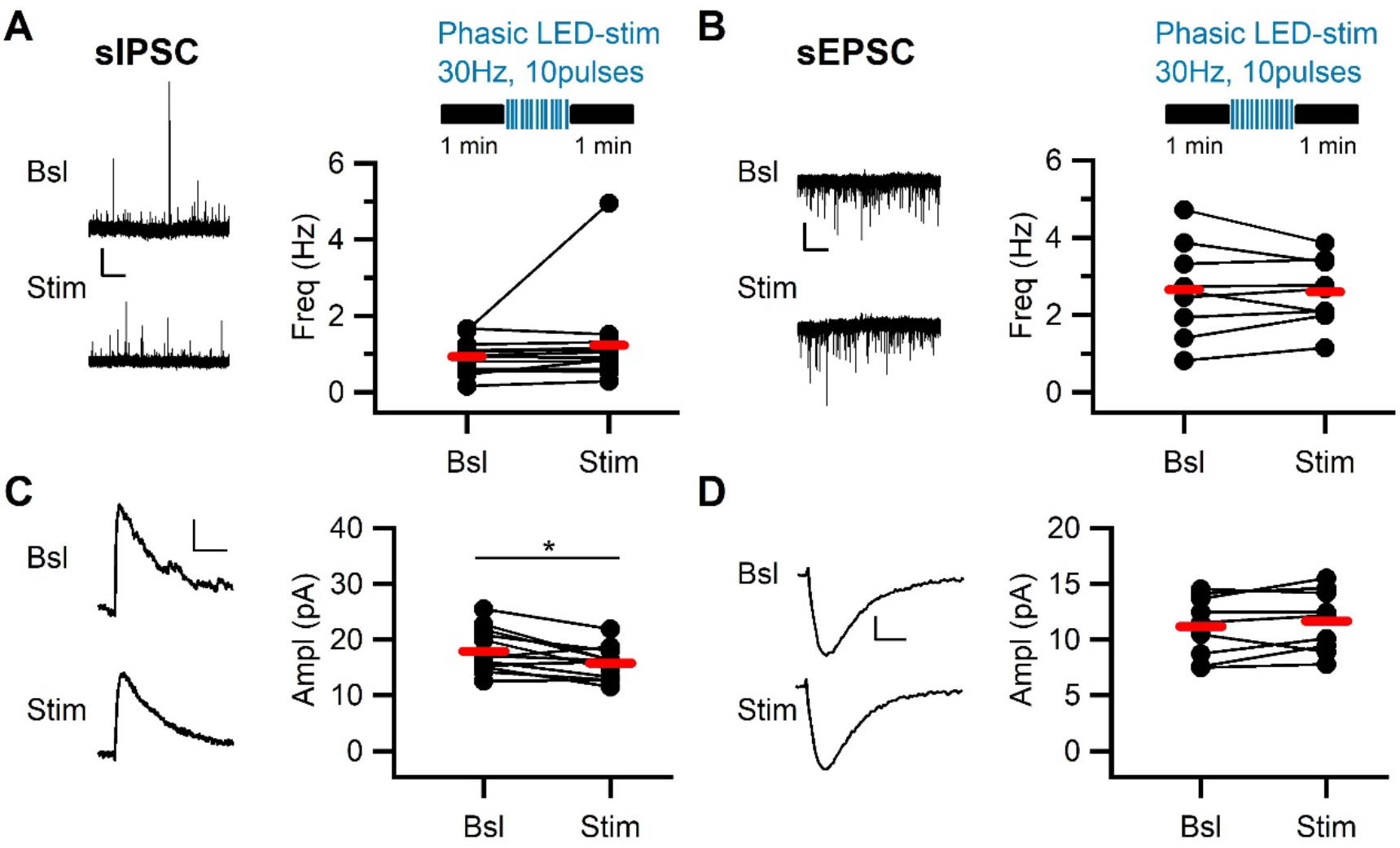
Phasic stimulation of midbrain dopaminergic fibers alters sIPSC amplitude in dm-ITCs. **(A and B)** Top: Experimental protocol. sIPSCs and sEPSCs were assessed before and after phasic stimulation of dopaminergic fibers (30 Hz, 10 pulses, 10 sweeps). Left: Example traces of sIPSC activitiy recorded at 0 mV (A) and sEPSC activity recorded at −70 mV in 100 μM PTX (B) recorded from dm-ITCs before (Bsl) and after stimulation (Stim). Scale bars: 20 pA, 10s. Right: Plot of sPSC frequency in individual neurons (dots) and average (red lines). sIPSC frequency (0.94 ± 0.12 Hz vs. 1.23 ± 0.32 Hz, n=13, paired t-test p=0.278) and sEPSC frequency (2.65 ± 0.40 Hz vs 2.60 ± 0.29 Hz, n=9, paired t-test p= 0.759) were not affected by dopaminergic fiber stimulation. **(C and D)** Left: Example traces of averaged sIPSCs (C) and sEPSCs (D) before and after stimulation. Scale bars: for sIPSC 5 pA, 20 ms, for sEPSC 3 pA, 3 ms. Right: Plot of sPSC amplitudes in individual neurons (dots) and average (red lines). sIPSC amplitude decreased after stimulation (17.94 ± 1.06 pA vs. 15.74 ± 0.78 pA, n=13, paired t-test *p=0.011), whereas sEPSC amplitude remained stable (11.18 ± 0.92 pA vs. 11.68 ± 0.92 pA; n=9, paired t-test p=0.217).

### DA modulates inhibitory synaptic transmission between ITC clusters

Following our observation that phasic stimulation of midbrain dopaminergic afferents affects sIPSCs, we wanted to more directly dissect the effect of DA on defined inhibitory inputs. Local inhibitory interactions have been shown between ITCs within one cluster, but also between medially located ITC clusters (Busti et al., 2011; Geracitano et al., 2007; Royer et al., 2000). To examine the effect of DA on between-cluster interactions, we used FoxP2-Cre mice to express AAV-ChR2-YFP specifically and selectively in one of the ITC clusters. In subsequent slice recordings, we stimulated axons and recorded in the innervated target cluster (Figure 4A, D, G). For the dm-ITCs, we hypothesized that the lateral ITC (l-ITC) cluster provides one of the inputs. Indeed, optical stimulation of ChR2-YFP+ fibers arising from l-ITCs reliably evoked PSCs in dm-ITCs (Figure 4B), that reversed around the equilibrium potential for chloride (Erev=−44.77 ± 1.77, n=6), and were largely abolished by the GABA_A_-channel blocker picrotoxin (PTX, % remaining current at 0 mV= 4.70 ± 1.09%, n=3). This suggests that l-ITCs provide fast GABAergic inputs to the dm-ITC cluster. Next, we assessed the effect of DA on inhibitory transmission in l-ITC→dm-ITC (Figure 4D) and dm-ITC→vm-ITC (Figure 4G) pathways. To select for effects of dopaminergic modulation of IPSCs without interference by GABA release from dopaminergic fibers, we opted to bath apply DA for 5 minutes. To gain insight into a possible presynaptic action, we used a paired pulse stimulation protocol (100 ms interstimulus interval). DA application significantly and reversibly suppressed IPSC amplitude to approximately 50% of baseline in l-ITC→dm-ITC and dm-ITC→vm-ITC pathways (Figure 4E and H), that was accompanied by a significant increase in the paired pulse ratio (PPR) (Figure 4F and I). Optically evoked IPSCs in the l-ITC→dm-ITC and dm-ITC→vm-ITC pathways were both of short latency, in keeping with monosynaptic connectivity (Figure 4C). Taken together, our data are consistent with a presynaptic site of action and indicate that DA dampens inhibitory interactions between l-ITC→dm-ITC and dm-ITC→vm-ITC clusters by decreasing GABA release. This is further supported by the fact that the decrease in IPSC amplitude was significantly correlated with the associated increase in PPR (Figure 5G).

**Figure 4.**
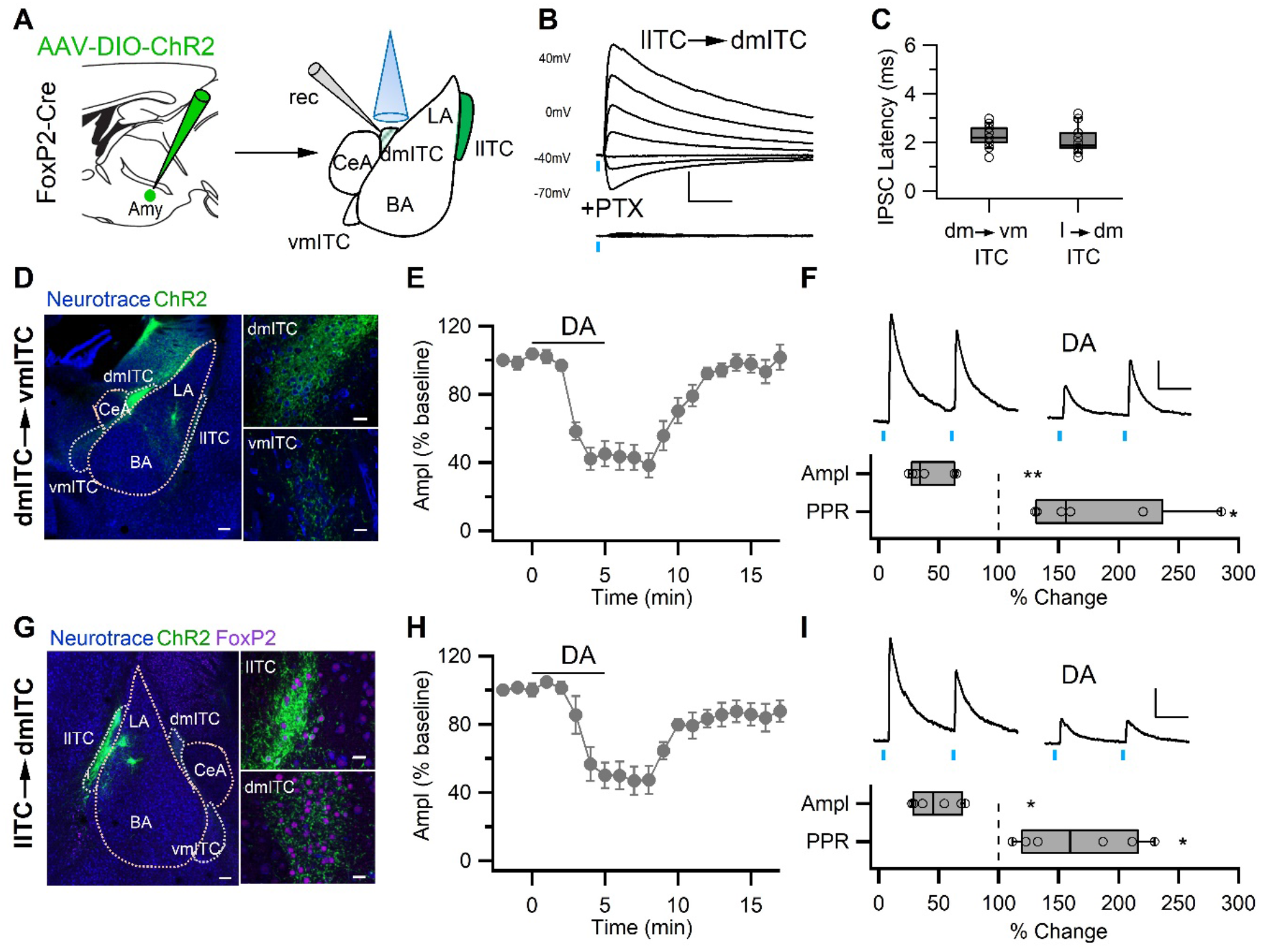
DA modulates inhibitory synaptic transmission between ITC clusters by a presynaptic mechanism. **(A)** Experimental scheme. AAV-DIO-ChR2-YFP was injected into the amygdala of FoxP2-Cre mice to specifically infect one ITC cluster. Upon targeting the l-ITC cluster, optically evoked PSCs were recorded in dm-ITCs. **(B)** PSCs recorded in a dm-ITC (l-ITC→dm-ITC pathway) at different holding potentials (−70 mV to +40 mV) with a reversal potential of −41.44 mV. Application of the GABA_A_ channel blocker PTX (100 μM) abolished the PSCs. Scale bars: 200 pA, 20 ms. **(C)** Box plots and individual latencies (circles) for IPSCs evoked between different ITC clusters. Latencies were similar for the l-ITC→dm-ITC pathway (2.07 ± 0.10 ms, n=22) and the dm-ITC→vm-ITC pathway (2.23 ± 0.09 ms, n=18, unpaired t-test p=0.268). **(D and G)** Confocal images of the amygdala showing ChR2-YFP expressing neurons and their efferents targeting dm-ITC (D) or l-ITC clusters (G). Smaller panels show the injection sites with YFP+ cells (top) and recording sites with YFP+ fibers (bottom). Scale bars: 200 μm and 20 μm. **(E and H)** Time course of changes in IPSC amplitude upon bath application of DA (30 μM, 5 min) in dm-ITC→vm-ITC and l-ITC→dm-ITC pathways. DA decreased IPSC amplitude, which returned to near baseline levels upon washout. **(F and I)** Top: Representative IPSC traces recorded at 0 mV from the baseline period and during DA application (paired pulse interval 100 ms). Bottom: Relative change of IPSC amplitude and paired pulse ratio (PPR) during DA application. Scale bars: 50 pA, 50 ms. **(F)** dm-ITC→vm-ITC pathway (n=6, amplitude 41.53 ± 7.25%, paired t-test **p=0.003; PPR 180.00 ± 25.01%, paired t-test *p=0.031). **(I)** l-ITC→dm-ITC pathway (n=6, amplitude 48.05 ± 8.09%, paired t-test **p= 0.006; PPR 165.72 ± 20.46%, paired t-test *p=0.019).

**Figure 5.**
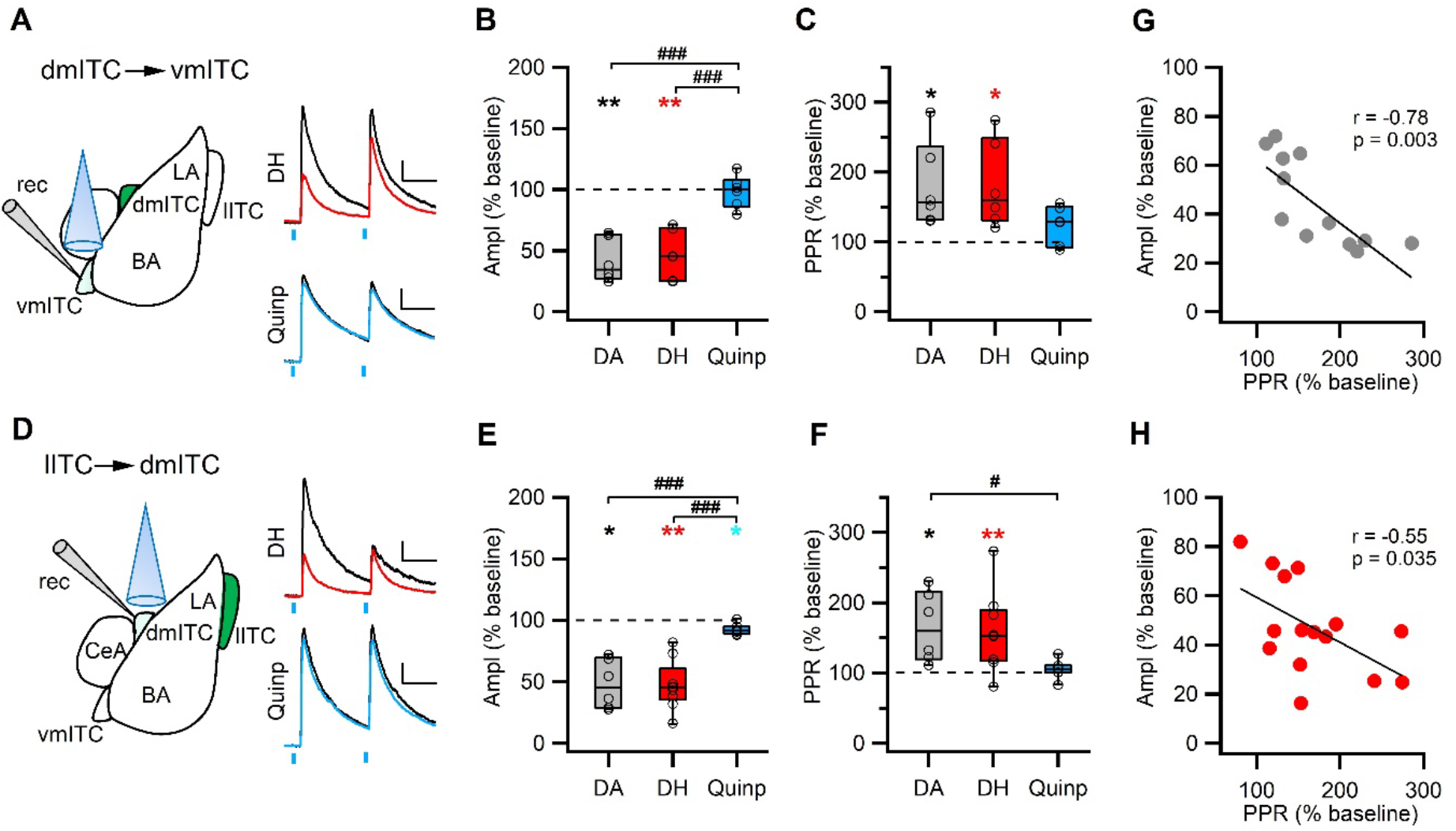
DRD1 mediates presynaptic depression between ITC clusters. **(A and D)** Schematic of experimental approach with infection, recording and light stimulation sites (left). Representative traces recorded at baseline (black) and during agonist application (colored). DH (10 μM), a DRD1 agonist, is shown in red, Quinpirole (Quinp, 1 μM), a DRD2 agonist, is shown in blue. Scale bars: 50 pA, 50 ms; paired pulse interval: 100 ms. **(B and E)** Comparison of relative changes in IPSC amplitude during agonist application in the dm-ITC→vm-ITC pathway (B) and l-ITC→dm-ITC pathway (E). **(B)** DH (red, n=6) suppressed IPSCs (amplitude 46.69 ± 8.14%, **p=0.010, paired t-test), whereas Quinp (blue, n=6) had no effect (amplitude 98.45 ± 5.36, p=0.438, paired t-test). Between group analysis revealed a significant drug effect on the dm-ITC→vm-ITC amplitude (one-way ANOVA, F (2, 15) =20.107, p<0.001) with Quinp differing from DA and DH (###p<0.001 each, Bonferroni post hoc tests). **(E)** DH (red, n=9) strongly suppressed IPSCs (amplitude 47.16 ± 6.67%, **p=0.003, paired t-test), whereas Quinp (blue, n=7) had a minor effect (amplitude 92.38 ± 1.63%, *p=0.017). Between group analysis revealed a significant drug effect on the l-ITC→dm-ITC amplitude (one-way ANOVA, F (2, 19) =17.394, p<0.001) with Quinp differing from DA and DH (###p<0.001 each, Bonferroni post hoc tests). **(C and F)** Comparison of relative change of PPR during drug application in the dm-ITC→vm-ITC pathway (C) and l-ITC→dm-ITC pathway (F). **(C)** DH (red, n=6) increased PPR (181.37 ± 25.38%, *p=0.035, paired t-test), whereas Quinp (blue, n=6) had no significant effect on PPR (123.97 ± 11.24%, p=0.127, paired t-test). Between group analysis revealed no significant drug effect on the dm-ITC→vm-ITC PPR (one-way ANOVA, F (2, 15) =2.304, p=0.134). **(F)** DH (red, n=9) increased PPR (158.46 ± 18.52%, **p=0.003, paired t-test), whereas Quinp (blue, n=7) had no effect on PPR (105.02 ± 4.98%, p=0.084, paired t-test). Between group analysis revealed a significant drug effect on the l-ITC→dm-ITC PPR (one-way ANOVA, F (2, 19) = 3.804, p=0.041). **(G and H)** Combined data from both pathways show a significant correlation between change in PPR and change in amplitude upon application of DA (G: Pearson correlation, r=−0.78, p=0.003, n=12) and upon application of DH (H: Pearson correlation, r=−0.55, p=0.035, n=15).

### Presynaptic DRD1 activation mimics the effect of DA in suppressing inhibitory interactions between ITC clusters

Dopamine receptors DRD1 and DRD2 have been localized in the amygdala, with ITC clusters being particularly enriched in DRD1 (Fuxe et al., 2003; Pinto and Sesack, 2008). To identify which receptor is involved in the DA-induced suppression of inhibitory synaptic transmission between ITC clusters, we used DH and quinpirole as selective agonists for DRD1 and DRD2, respectively.

Activation of DRD1 significantly suppressed IPSCs in the l-ITC→dm-ITC pathway, whereas DRD2 activation had no effect (Figure 5A-B). Furthermore, activation of DRD1 significantly increased the PPR in the l-ITC→dm-ITC pathway, whereas DRD2 activation did not (Figure 5A and C). The DH-induced IPSC suppression and PPR increase was comparable to the effect of DA application (Figure 5B-C). Likewise, DRD1 activation also suppressed IPSCs similar to DA (to about 50%) in the dm-ITC→vm-ITC pathway, whereas DRD2 activation had only a small, yet significant effect on amplitude, that was different from that of DA or DRD1 activation (Figure 5D-E). Activation of DRD1 also significantly increased the PPR in the dm-ITC→vm-ITC pathway, but DRD2 activation did not (Figure 5E-F). Moreover, we again observed a significant correlation between the decrease in IPSC amplitude and the increase in PPR upon application of the DRD1 agonist (Figure 5H), supporting a presynaptic site of action.

Another reported mechanism by which DA tunes ITC output in younger animals is a direct DRD1-mediated hyperpolarization (Manko et al., 2011; Marowsky et al., 2005). Therefore, we also tested for a direct postsynaptic action of DA by recording dm-ITC neurons in current-clamp mode from young and adult GAD67-GFP mice without stimulating any input (Figure 5–figure supplement 1A). We observed a significant DA-induced hyperpolarization in about 50% of recorded dm-ITC cells from both age groups (Figure 5–figure supplement 1B-D). Thus, irrespective of age, only a fraction of ITC neurons exhibits a small, direct postsynaptic response to application of DA. In conclusion, our data suggests that DA modulates ITC cluster interactions chiefly by presynaptic DRD1 receptors.

### Early extinction training alters midbrain input properties and modulation of ITCs

Recent data show that activity in midbrain dopaminergic neurons correlates strongly with early extinction learning, suggesting that in the amygdala, DA provides a prediction error-like neuronal signal that is necessary to initiate fear extinction (Luo et al., 2018; Salinas-Hernandez et al., 2018). Thus, we tested if early extinction training alters co-release from midbrain dopaminergic inputs to ITC neurons and the dopaminergic modulation of ITC cluster interaction. To this end, we subjected one group of mice to fear conditioning on day 1 and a day later to fear retrieval and early extinction training (E-Ext) by presenting 16 CSs in the absence of the reinforcing US, while the control group was only exposed to the same number of CSs (CS-only, Figure 6A). In both DAT-IRES-Cre and FoxP2-Cre mice transduced with ChR2-YFP in midbrain and ITC neurons, respectively, this short protocol did not reduce fear expression significantly (Figure 6–figure supplement 1A-B and figure supplement 3A-B). Importantly, these mice can extinguish when subjected to a 2-day extinction training protocol (Figure 6–figure supplement 2). As our focus was on early extinction, we opted to obtained acute brain slices for recordings from ChR2-YFP transduced mice only after early extinction training (Figure 6A).

**Figure 6.**
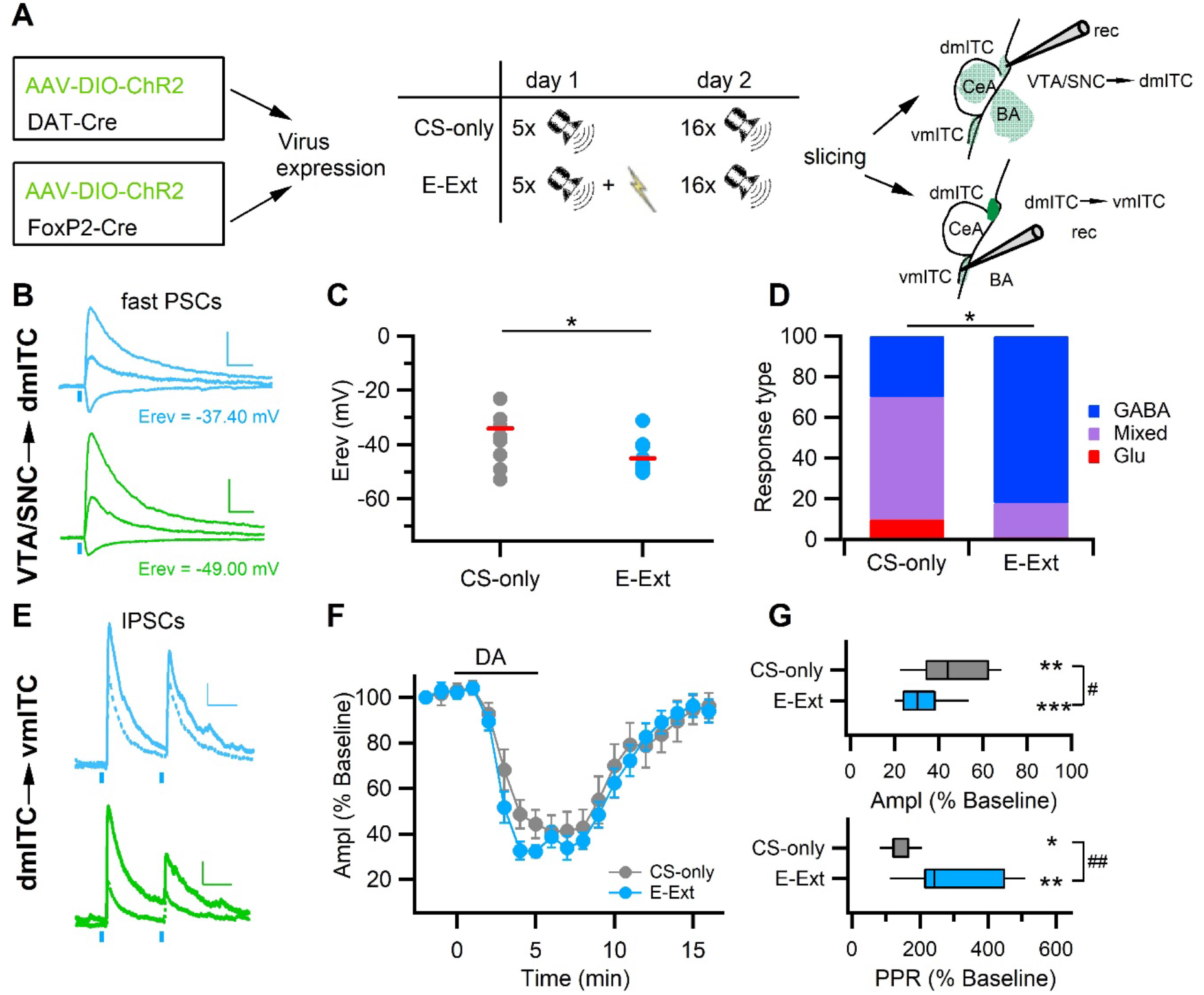
Early extinction enhances GABA release from midbrain terminals and DA-mediated depression of the dm-ITC→vm-ITC pathway. **(A)** Experimental scheme: To investigate the VTA/SNC→dm-ITC pathway, ChR2 was transduced in dopaminergic midbrain neurons of DAT-IRES-Cre mice. To investigate DA modulation of the dm-ITC→vm-ITC pathway, ChR2 was transduced into the dm-ITC cluster of FoxP2-Cre mice. The early extinction group (E-Ext) underwent fear conditioning on day 1 (5 CS-US pairings) and early extinction training on day 2 (16 CS presentations). The CS-only group received only CS presentations. **(B)** Left: Example traces of light-evoked PSCs by dopaminergic fiber stimulation recorded in dm-ITCs at −70mV, 0mV, and 40mV from CS-only (grey traces) or E-Ext animals (blue traces). Scale bars: 50 pA, 50 ms. **(C)** Plot of PSC reversal potentials in individual dm-ITCs (dots) and average (red lines) from CS-only and E-Ext groups. Erev was significantly lower in the E-Ext (−45.09 ± 1.76 mV, n=10 cells from 6 animals) vs. the CS-only group (−34.06 ± 5.11 mV n=11 cells, from 4 animals, *p=0.047, paired t-test). **(D)** Summary graph comparing the type of fast PSCs in dm-ITCs recorded from CS-only and E-Ext animals (CS-only, n=10: GABA 30%, mixed 60%, Glu 10%; E-Ext, n=11: GABA 82%, mixed 18%). PSC types were significantly different between groups (Fisher’s exact test =5.68, *p=0.041). **(E)** Example traces of light-evoked IPSCs recorded in vm-ITCs at 0 mV upon paired pulse stimulation (100 ms interstimulus interval) of the dm-ITC→vm-ITC pathway from CS-only (black traces) or E-Ext (blue traces) animals before (solid) and during DA application (dotted). Scale bars: 50 pA, 50 ms. **(F)** Time course of changes in IPSC amplitude upon bath application of DA (30 μM, 5 min) in dm-ITC→vm-ITC pathway in CS-only and E-Ext groups. Two-way ANOVA (1-9 min) revealed significant changes for time F(8)=37.903, p<0.001 and group F(1)=8.229, p = 0.005 but no significant interaction F(8, 144)=0.521, p=0.839. **(G)** Significant changes of IPSC amplitude (paired t-tests; CS-only, **p=0.002, CS-US ***p<0.001) and PPR (paired t-tests; CS-only, *p=0.035, CS-US **p=0.003) 4-5 min after DA application in both groups. IPSC amplitude was more depressed in neurons recorded from E-Ext vs. CS-only animals (32.42 ± 3.15%, n=11 cells from 6 animals, vs. 46.45 ± 6.00%, n=7 cells from 4 animals, unpaired t-test #p=0.036). The PPR increase was larger in neurons recorded from E-Ext. vs. CS-only animals (310.63 ± 44.63%, n=11, vs. 140.649 ± 15.22%, n=7, unpaired t-test ##p=0.007).

Early extinction training significantly decreased the reversal potential of fast PSCs evoked by stimulation of dopaminergic inputs from confirmed midbrain injection sites (Figure 6–figure supplement 1C) onto dm-ITCs compared to animals in the CS-only group (Figure 6B-C). When categorizing PSCs according to their reversal potential, as described for naïve animals (c.f. Figure 2C-F), we found that the pattern of responses shifted to more GABAergic PSCs with early extinction training (Figure 6D). This suggests that midbrain inputs are more likely to co-release GABA, that can contribute to inhibit dm-ITCs. Secondly, we found that early extinction training altered the efficacy of dopaminergic modulation of ITC cluster interaction in the dm-ITC→vm-ITC pathway (Figure 6E-G). The suppression of IPSC amplitude by DA was significantly larger in neurons recorded from animals in the E-Ext vs. CS-only group, with a concomitant significantly larger increase in the PPR of IPSCs (Figure 6F-G). This suggests that early extinction training decreases inhibition by enhancing a dopaminergic presynaptic mechanism, which can contribute to the disinhibition of vm-ITCs. In summary, our data indicate that changes in dopaminergic input action during early extinction learning can switch the activity balance between ITC clusters by enhancing inhibition of dm-ITCs and disinhibition of vm-ITCs.

## Discussion

Here, we investigated the functional impact of midbrain dopaminergic inputs and dopaminergic modulation onto amygdala ITCs. Our key findings are that ITCs are controlled by two distinct mechanisms including fast inhibition resulting from a prominent co-release of GABA, and a slower DRD1-mediated presynaptic suppression of inhibitory interactions between distinct ITC clusters (Figure 7A). Upon early extinction learning, fast inhibition onto dm-ITCs is increased, and DA more potently suppresses dm cluster-mediated inhibition of vm-ITCs. This may support a shift in the activity balance between these two distinct ITC clusters by inhibiting dm-ITCs and disinhibiting vm-ITCs to enable fear suppression during extinction learning (Figure 7B). Although it is well established that amygdala ITC clusters receive dense TH+ projections (Asan, 1997; Fuxe et al., 2003), here, we show that these are part of the mesolimbic pathway from VTA/SNC, also providing afferents to amygdalostriatal transition zone (Astria), BA, and to a lesser extent to CeA. Our results are well in line with a recent report that 6-hydroxy-dopamine injections in BLA resulted in the loss of TH+ neurons in VTA/SNC that also denervated ITCs and ventral Astria, but not CeA (Ferrazzo et al., 2019). The CeA was shown to receive largely distinct dopaminergic inputs arising from dorsotegmental nuclei (Groessl et al., 2018). Consistent with previous ultrastructural investigations demonstrating that TH+ axons contact ITC somata, dendrites, and spines (Asan, 1997; Pinto and Sesack, 2008), we detected putative presynaptic terminals originating from VTA/SNC on the soma and along ITC dendrites suggesting functional connectivity.

**Figure 7.**
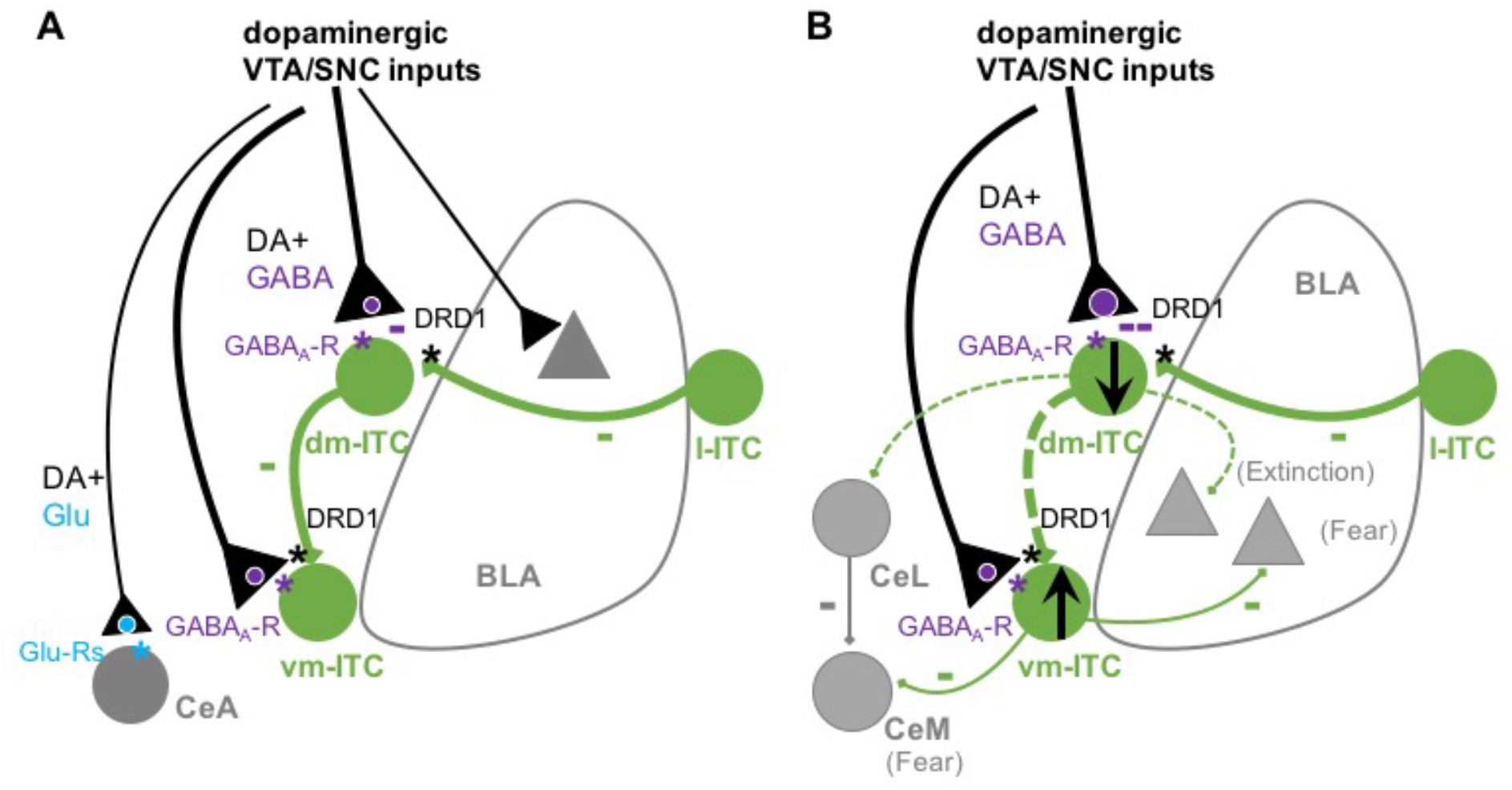
Summary scheme of results. **(A)** Dopaminergic inputs from VTA/SNC target amygdala ITCs and to a lesser extent BLA and CeA. Co-release of glutamate is mainly observed in CeA, whereas GABA co-release is prominent onto dm- and vm-ITCs. DA depresses inhibitory interactions between ITCs clusters via presynaptic DRD1. Activation of midbrain inputs can thus regulate the ITC network by rapid inhibitory and slower disinhibitory mechanisms. **(B)** Early extinction training enhances both inhibition of dm-ITCs, by biasing towards GABA co-release, and disinhibiton of vm-ITCs, by altering DRD1-mediated suppression of the dm-ITC→vm-ITC pathway. Together, this may tip the activity balance towards decreased dm-ITC and increased vm-ITC activity. We speculate that this could impact behavioral outcome by decreasing inhibition onto dm-ITC targets (such as CeL and extinction-promoting neurons in BLA), while promoting inhibition onto vm-ITC targets (such as fear-promoting neurons in CeM and BLA).

While co-release of glutamate and GABA from VTA/SNC dopaminergic neurons is well established in nucleus accumbens and dorsal striatum (Granger et al., 2017), glutamate co-release has only recently been demonstrated in CeA (Mingote et al., 2015). Here, we also find GABA co-release in CeA, although to a much lower extent than glutamate co-release. We did not observe glutamate co-release in the BLA, which is also in line with a previous study (Mingote et al., 2015), but see also (Lutas et al., 2019).

Our study is the first to directly examine dopaminergic afferents onto amygdala ITCs via optogenetic stimulation. Combined qualitative anatomical and quantitative electrophysiological evidence strongly supports the notion that glutamate and GABA can be released from dopaminergic afferents onto ITCs, with a major contribution of GABA. GABA co-release is a prominent feature of midbrain dopaminergic neurons targeting dorsal striatum and nucleus accumbens, where it can rapidly inhibit medium spiny neurons of the direct and indirect pathways (Tritsch et al., 2012). The presence of vGluT1/2 and vGAT in dopaminergic fibers innervating medial ITC clusters is in accordance with canonical release mechanisms for glutamate and GABA, respectively. In dorsal striatum however, GABA is transported into vesicles via the vesicular monoamine transporter VMAT2 suggesting a spatio-temporal synchronization of GABA and DA release from the same vesicles (Tritsch et al., 2012). Which of the two mechanisms is predominant in ITC afferents remains to be investigated. Furthermore, in dopaminergic striatal afferents, GABA is either synthesized in a non-canonical aldehyde-dehydrogenase1a1-dependent pathway, and/or relies on GABA uptake from the extracellular milieu via GABA transporters (Kim et al., 2015; Tritsch et al., 2014). Although we did not determine which mechanism provides GABA for inhibiting ITCs, regulation of GABA uptake could endow midbrain afferents with the flexibly to rapidly alter GABAergic co-transmission in a target-specific manner. It may, therefore, be a good candidate mechanism mediating changes in GABA co-release during early extinction learning.

ITCs receive a number of well characterized excitatory inputs from BLA, thalamus, prefrontal and sensory cortex that drive their activity (Amir et al., 2011; Asede et al., 2015; Cho et al., 2013; Pare et al., 2003; Strobel et al., 2015). Apart from local inhibitory interactions between ITCs, that were proposed to stabilize overall spike output (Geracitano et al., 2007; Manko et al., 2011; Royer et al., 2000), it remained unclear which extrinsic inputs inhibit ITCs. Co-release of GABA from dopaminergic midbrain could provide a rapid and temporally precise signal to inhibit ITC spike activity in response to salient stimuli and/or during learning. Furthermore, spatially targeted inhibition onto ITC dendrites or spines may shunt glutamatergic inputs to enable local control of synaptic plasticity (Tritsch et al., 2016).

Phasic activity of midbrain dopaminergic neurons may signal unexpected rewards or aversive events such as noxious stimuli and these activity patterns shape DA release in target structures (Brischoux et al., 2009; Bromberg-Martin et al., 2010). Phasic stimulation (> 20 Hz) of VTA neurons increases DA release in forebrain structures, including amygdala, and affects behavior (Holloway et al., 2018; Tsai et al., 2009). In our hands, phasic but not tonic stimulation of midbrain dopaminergic inputs to dm-ITCs depressed sIPSC, suggesting that released DA modulates inhibitory transmission. Our results are in agreement with a recent report showing that DA depressed IPSC amplitude in vm-ITCs (Gregoriou et al., 2019). Despite earlier studies reporting that DA directly hyperpolarizes medial and lateral ITCs via postsynaptic DRD1 (Manko et al., 2011; Marowsky et al., 2005), Kwon and colleagues did not replicate these findings (Kwon et al., 2015). Our data may resolve this discrepancy by demonstrating that only a fraction of dm-ITCs is directly hyperpolarized by DA, and indicate the presence of several distinct dopaminergic mechanisms gating ITC activity.

While ITCs form local inhibitory networks within, but also between medially located neighboring clusters (Amir et al., 2011; Busti et al., 2011; Geracitano et al., 2007; Manko et al., 2011; Royer et al., 2000), our novel finding that l-ITCs provide input to dm-ITCs suggests further complexity of inhibitory and disinhibitory interactions between ITC clusters that can provide a substrate for integration of information from distinct afferents (Asede et al., 2015; Morozov et al., 2011; Strobel et al., 2015). DA also modulated the interactions between ITC clusters. Its disinhibitory effect via presynaptic DRD1 in the target cluster resembles the situation in the nucleus accumbens, where presynaptic DRD1 attenuates IPSCs and consequently, lateral inhibition between medium spiny neurons, a mechanism proposed to diminish competitive interactions between single projection neurons or ensembles (Nicola and Malenka, 1997; Pennartz et al., 1992; Taverna et al., 2005). In analogy, the same mechanism may enable selection of specific ITC clusters or ensembles during distinct behavioral states. Our data extend current views on dopaminergic modulation of the ITC network with DRD1 playing a major role both pre- and postsynaptically. While direct inhibition of ITCs decreases overall inhibition onto diverse downstream regions, target-specific presynaptic disinhibition would promote inhibition from distinct ITC clusters onto defined downstream regions. These mechanisms may help to sharpen competitive interactions between clusters and to select a defined ITC network output. In the context of behavior, in vivo pharmacological studies in amygdala pointed to a role of DRD1 in the acquisition of cued fear memory (Guarraci et al., 1999; Lamont and Kokkinidis, 1998; Nader and LeDoux, 1999) as well acquisition of extinction (Hikind and Maroun, 2008). Cellular effects of DRD1 include altered excitability of BLA projection- and local interneurons, as well as DRD1-dependent LTP in the BLA-CeA pathway that have been discussed in the context of fear acquisition (Groessl et al., 2018; Lee et al., 2017). However, DRD1-dependent ITC network modulation could provide the required flexibility to support behavioral transitions during acquisition of extinction memory. Circuit and immediate early gene mapping studies converged on a model where dm-ITC become engaged and undergo plasticity during fear learning keeping vm-ITC activity in check. During extinction learning, this balance tips towards recruitment of vm-ITCs, which are required for extinction retrieval (Busti et al., 2011; Duvarci and Pare, 2014; Likhtik et al., 2008). Our data suggest that dopaminergic afferent control over ITCs could play a critical role in these transitions. DA neurons in the VTA are activated by unexpected omission of the US implying that they provide a prediction-like error signal to downstream targets including ITCs (Luo et al., 2018; Salinas-Hernandez et al., 2018). Consistent with the above model, enhanced GABA co-release onto dm-ITCs during early extinction could rapidly decrease their activity, and together with enhanced DRD1-mediated disinhibition of vm-ITCs allow for activation of the latter. Downstream, dm- and vm-ITCs most likely exert behavioral output by differentially inhibiting specific fear- and extinction related subpopulations of BLA and CeL or centro-medial (CeM) neurons (Figure 7B) (Asede et al., 2015; Duvarci and Pare, 2014; Gregoriou et al., 2019). Concurring or subsequent synaptic plasticity at BLA and other afferents onto dm- and vm-ITCs could help maintain this activation pattern during extinction retrieval (Amano et al., 2010; Asede et al., 2015; Huang et al., 2014; Kwon et al., 2015).

While our study delineates several novel cellular mechanisms how mesolimbic dopaminergic afferents control amygdala ITC networks, the observed *ex vivo* alterations are still correlative. Further studies are warranted to understand how these cellular mechanisms impact fear and extinction learning and memory. Beyond fear memories, DA is critical for processes related to incentive salience, motivation, and cue-reward learning (Abraham et al., 2014; Bromberg-Martin et al., 2010). For example, VTA dopaminergic inputs to BLA are activated by motivationally salient appetitive and aversive outcomes and acquire responses to predictive cues during learning (Lutas et al., 2019). DA is also released in the amygdala during affective states such as stress (Belujon and Grace, 2015; Yokoyama et al., 2005). Therefore, it is intriguing to speculate that dopaminergic modulation of distinct ITCs clusters may also play a role in other processes such as reward learning and addiction, as well as in the control of mood-related behavior such as anxiety- and depression-like behavior (Ferrazzo et al., 2019; Kuerbitz et al., 2018).

## Materials and Methods

### Animals

We used 6 to 14 week-old adult male DAT-IRES-Cre (JAX#006660, Jackson Laboratories, Bar Harbor, Maine, USA) (Backman et al., 2006) or FoxP2-Cre (JAX#030541, Jackson Laboratories) (Rousso et al., 2016) transgenic mice for Cre-dependent expression of viral vectors, and 20-28 day-old (young) and 8-10 week-old adult male GAD67–GFP (Tamamaki et al., 2003) mice for slice experiments without optical stimulation. All lines were heterozygous and backcrossed to C57BL6/J. Mice were kept in a 12-hour light/dark (6 am to 6 pm) cycle with access to food and water ad libitum. All behavioral experiments were conducted during the light cycle (between 7 and 10 am). All animal procedures were performed in accordance with institutional guidelines and with current European Union guidelines, and were approved by the local government authorities for Animal Care and Use (Regierungspraesidium Tuebingen, State of Baden-Wuerttemberg, Germany).

### Surgical procedures

Stereotaxic injection of AAV-EF1a-DIO-HChR2(H134R)-eYFP (serotype 2/1 or 2/9, U. Penn Vector Core, Philadelphia, PA, USA or Addgene, Watertown, MA, USA) was performed in 6-8 weeks old DAT-IRES-Cre or FoxP2-Cre transgenic mice. Mice were anesthetized with isoflurane in oxygen-enriched air (Oxymat 3, Weinmann Medical Technologies, Hamburg, Germany) and the head was fixed in a stereotaxic frame (Kopf Instruments, Tujunga, CA, USA or Stoelting, Wood Dale, IL, USA). Eyes were protected with ointment and the body temperature of the animal was maintained using a feedback-controlled heating pad with a rectal sensor (FHC, Bowdoin, ME, USA).

Lidocaine was used as a local anesthetic. An incision was made on the skin, the skull was exposed and coordinates of bregma were identified. The skull was drilled with a microdrill (Kopf Instruments, Tujunga, CA, USA) at the desired coordinates in reference to bregma. Borosilicate capillaries for VTA/SNC (1B150F-4, World Precision Instruments, Friedberg, Germany) and ITC injections (marked 1 to 5 μl, Drummond Scientific, Broomall, PA, USA) were pulled on a horizontal pipette puller (P-1000, Sutter Instruments, Novato, CA, USA) and used to pressure inject viruses at a volume of 300-500 nl for VTA/SNC and 25–50 nl for ITCs. The capillaries were slowly withdrawn, the skull disinfected and the skin sutured with silk. Postoperative pain medication included injection of meloxicam (Metacam, Boehringer Ingelheim, Ingelheim, Germany) at 5 mg/kg subcutaneously. The following coordinates were used in reference to bregma: for VTA/SNC in mm, AP: −3.00, ML: ±0.50, DV: 4.50; for dm-ITC in mm, AP: −1.40, ML: ±3.3, DV: 4.70; for l-ITC in mm, AP: −1.40, ML: ±3.45, DV: 4.70).

### Slice recordings and analysis

Three (for ITCs) to six (for VTA/SNC) weeks after viral injections, mice were deeply anaesthetized with 3% isoflurane (Isofluran CP, cp-pharma, Burgdorf, Germany) in oxygen and decapitated. The brain was rapidly extracted and cooled down in ice-cold slicing artificial cerebrospinal fluid (ACSF) containing (in mM): 124 NaCl, 2.7 KCl, 26 NaH2CO3, 1.25 NaH2PO4, 10 MgSO4, 2 CaCl2, 18 D-Glucose, 4 ascorbic acid, equilibrated with carbogen (95% O_2_/5% CO_2_). Coronal brain slices (320 μm) containing the amygdala were cut in ice-cold slicing ACSF with a sapphire blade (Delaware Diamond Knives, Wilmington, DE, USA) on a vibrating microtome (Microm HM650V, Thermo Scientific, Dreieich, Germany). Slices were collected in a custom-built interface chamber with recording ACSF containing (in mM): 124 NaCl, 2.7 KCl, 26 NaH_2_CO_3_, 1.25 NaH_2_PO_4_, 1.3 MgSO_4_, 2 CaCl_2_, 18 D-Glucose, 4 ascorbic acid, equilibrated with carbogen. Slices were recovered at 37°C for 40 minutes and stored at room temperature. Whole-cell patch-clamp recordings were performed in a submersion chamber under an upright microscope (Olympus BX51WI, Olympus Germany, Hamburg, Germany), where slices were superfused with recording ACSF at 30-31°C. Recordings were performed using an Axon Instruments Multiclamp 700B amplifier and a Digidata 1440A digitizer (both Molecular Devices, San Jose, CA, USA). Glass micropipettes (6-9 MΩ resistance when filled with internal solution) were pulled from borosilicate capillaries (ID 0.86 mm, OD 1.5 mm, Science Products, Hofheim, Germany).

Recordings in voltage-clamp configuration were performed with cesium-based internal solution containing (in mM): 115 Cs-methanesulphonate, 20 CsCl, 4 Mg-ATP, 0.4 Na-GTP, 10 Na_2_-phosphocreatine, 10 HEPES, 0.6 EGTA (290-295 mOsm, pH 7.2-7.3). Signals were low-pass filtered at 2 kHz and digitized at 5 kHz. Recordings in current-clamp configuration were performed with K-Gluconate based internal solution containing (in mM): 130 K-Gluconate, 5 KCl, 4 Mg-ATP, 0.4 Na-GTP, 10 Na2-phosphocreatine, 10 HEPES, 0.6 EGTA (290-295 mOsm, pH 7.2-7.3). Signals were low-pass filtered at 10 kHz and digitized at 20 kHz. Changes in series resistance <30% were accepted. Optical stimulation was achieved by triggering a light-emitting diode (LED; 470 nm, KSL70, Rapp Opto-Electronics, Hamburg, Germany or CoolLED pE, CoolLED, Andover, UK) coupled to the upright microscope objective (Olympus Germany, 60x 1.0 NA). PSC were recorded from LED stimulations (pulse length 0.2-5 msec) with 0.1 Hz frequency. Spontaneous and evoked IPSCs were recorded at 0 mV holding potential. For sEPSC recordings, GABAergic activity was blocked with PTX and cells were held at −70 mV. Phasic stimulation of dopaminergic fibers consisted of 10 pulses at 30 Hz repeated 10 times with 10 second inter-sweep interval. Tonic stimulation consisted of a single sweep of 100 pulses at 1 Hz. In some experiments biocytin (3-5%) was added to the internal solution for post-hoc visualization of the recorded cells.

Electrophysiological data were analyzed with Neuromatic (www.neuromatic.thinkrandom.com) (Rothman and Silver, 2018) and/or custom written functions in Igor Pro (Wavemetrics, Portland, OR, USA). PSC amplitudes were measured as the peak value in reference to a 5 msec baseline period. For co-release data, reversal potential (Erev) was calculated by regression analysis of the peak current at different holding potentials (−70 mV, −50 mV, 0 mV, +40 mV). Cells with Erev <-40 mV were considered to have GABAergic, −40 mV< Erev <-15 mV to have mixed, and Erev >-15mV to have glutamatergic PSCs.

Chemicals were purchased from Roth, Merck or Sigma. Drugs for pharmacology were obtained from Sigma, Tocris or Biotrend. All drugs were diluted in ACSF from concentrated frozen stocks on the day of recording. Drugs were bath-applied at a rate of 2 ml/minute using a peristaltic pump (Ismatec Products, Cole-Palmer, Wertheim, Germany).

### Behavioral procedures

Fear conditioning was performed in a square chamber with a grid floor that was cleaned with 70% ethanol (Context A). Five pairings of conditioned (CS) and unconditioned (US) stimuli were presented. The CS was a 30 second tone (at 7.5 kHz, 80 dB) which coincided at its offset with the US (1 second foot shock, 0.4 mA). In the CS-only group, the US was omitted. Fear extinction was performed in a round chamber with a flat floor that was cleaned with 1% acetic acid (Context B). Early extinction training started 24 hours after fear conditioning and consisted of 16 CSs. Full extinction training was performed by presenting 25 CSs 24 hours and 48 hours after conditioning. All sessions started with a two-minute baseline assessment and the subsequent CSs were presented at random intervals of 20 to 180 seconds. Freezing was detected using an infrared beam detection system (Coulbourn Instruments, Holliston, MA, USA), with the threshold for freezing set to 2 seconds of immobility and quantified offline with custom-written macros in Microsoft Excel (Asede et al., 2015). Freezing data for fear conditioning are shown as average post-shock freezing for the last two CS-US pairings. CS-induced freezing during extinction is shown as averages four CSs. Animals were randomly assigned to experimental groups. Electrophysiology experiments were performed 80-90 minutes after the behavioral procedures by an experimenter blind to the training procedure of the animal.

### Staining, immunohistochemistry, and imaging procedures

Slices containing midbrain and ITC injection sites were fixed in 4% paraformaldehyde (PFA) in phosphate buffered saline (PBS), whereas recorded slices with biocytin filled cells were fixed in 4% PFA, 15% saturated picric acid solution, and 0.05% glutaraldehyde in PBS overnight at 4°C. All slices were resectioned on a vibratome (Microm HM650V, Thermo Scientific, Dreieich, Germany) at 60 to 65 μm and processed as previously described (Asede et al., 2015) with minor modifications.

Biocytin was revealed using fluorescently-conjugated Streptavidin (1:1000, Dianova, Germany). Immunostainings were performed using standard procedures. Upon blocking with PBS complemented with 0.3% Triton and 10% serum for 90 minutes, the following primary antibodies were used: goat anti-GFP (GeneTex, 1:500), rabbit anti-GFP (Invitrogen, 1:750), rabbit anti-FoxP2 (Abcam, 1:1000), mouse anti-FoxP2 (Merck Millipore, 1:1000), rabbit anti-TH (Millipore, 1:1000), mouse anti-TH (Merck, 1:1000), mouse anti-Bassoon (Abcam, 1:500). Secondary antibodies used were: Alexa488-conjugated donkey-anti-rabbit, Alexa555-conjugated donkey-anti-rabbit and goat-anti-mouse, Alexa647-conjugated donkey-anti-rabbit, Alexa633-conjugated goat-anti-mouse and goat-anti-rabbit. All secondary antibodies were obtained from Invitrogen and used in 1:1000 dilutions.

Overview images were taken with a fluorescence microscope (Axio Imager, Carl Zeiss, Oberkochen, Germany). Detailed images of injection and projection sites were taken with a LSM710 laser-scanning microscope (Carl Zeiss, Oberkochen, Germany) equipped with either a 40x 1.3 NA or a 63x 1.4 NA objective, with the pinhole set to 1 airy unit. Most of the confocal images are presented as maximum intensity projection of z-stacks. For deconvolution, Huygens suite 19.04 (Scientific Volume Imaging, Hilversum, Netherlands) was used. Cell counts were performed from confocal z-stack images with the cell counter plugin FIJI in ImageJ (https://imagej.net/Cell_Counter). Specificity of the infections in the dopaminergic midbrain was calculated as the ratio of double labeled cells (TH+GFP+ cells) to all infected cells (GFP+ cells).

For analysis of presynaptic markers, 60 μm coronal sections containing the ITCs, from DAT-IRES-Cre mice injected with AAV-EF1a-DIO-HChR2(H134R)-eYFP, were immunostained for vGluT1/2 or vGAT. Sections were incubated with the following primary antibodies: polyclonal rabbit anti-GFP (Invitrogen 1: 1000), sheep anti-FoxP2 (RnDSystems, 1:200) and polyclonal guinea pig anti-vGAT (Synaptic Systems, 1:500) or a combination of guinea pig-anti-vGluT1 (Millipore, 1:3000) and vGluT2 (Chemicon International, 1:5000). Secondary antibodies used were: AlexaFluor 488 donkey anti rabbit (life technologies, 1:1000), Cy5 donkey anti goat (Jackson Immunoresearch Lab., 1:400) and DyLight 405 donkey anti guinea pig (Jackson Immunoresearch Lab., 1:500). Images of vGluT and vGAT stainings were visualized using an Airy Scan LSM980 laser scanning microscope (Carl Zeiss, Oberkochen, Germany) with a 40x 1.3 NA objective. The pinhole was set to 1 airy unit. Z-stacks were obtained and then analyzed in 3D using Imaris Software (Bitplane, Zurich, Switzerland).

### Data presentation and statistics

Most data are represented as individual points, box and whisker plots (reporting median, 25, 75, 10, and 90 percentiles), or bar graphs of average ± SEM. Where applicable, electrophysiology data were normalized to the first minute of baseline recording, similarly Z-scores were calculated from the average and standard deviation of the baseline.

No methods were used to predetermine sample size. Sample sizes are similar to other studies in the field. Statistical analysis was performed using the program SPSS (IBM, Germany). A p-value <0.05 was considered significant. Paired or unpaired Students t-tests were used for two dependent or independent comparisons, respectively. Comparison of multiple groups was done using one- or two-way ANOVA followed by a Bonferroni post-hoc test if appropriate. Repeated treatments were compared using repeated measure ANOVA followed by a Bonferroni post-hoc test if appropriate. Categorical data were compared with Fisher’s exact test.

## Acknowledgements

We thank Johannes Ungermann and Marlly Achury for help with histology, members of the Ehrlich and Ferraguti labs for discussion and support, and Dr. Julien Genty, Martin Zeller, and Prof. Norbert Hajos for critical reading and commenting on an earlier version of the manuscript. This work was supported by the Charitable Hertie Foundation and the German Research Foundation DFG EH 197/3-1 (to I.E.) and by the Austrian Science Fund (FWF) grant # I-2215 (to F.F).

---

**Figure 1–figure supplement 1.**
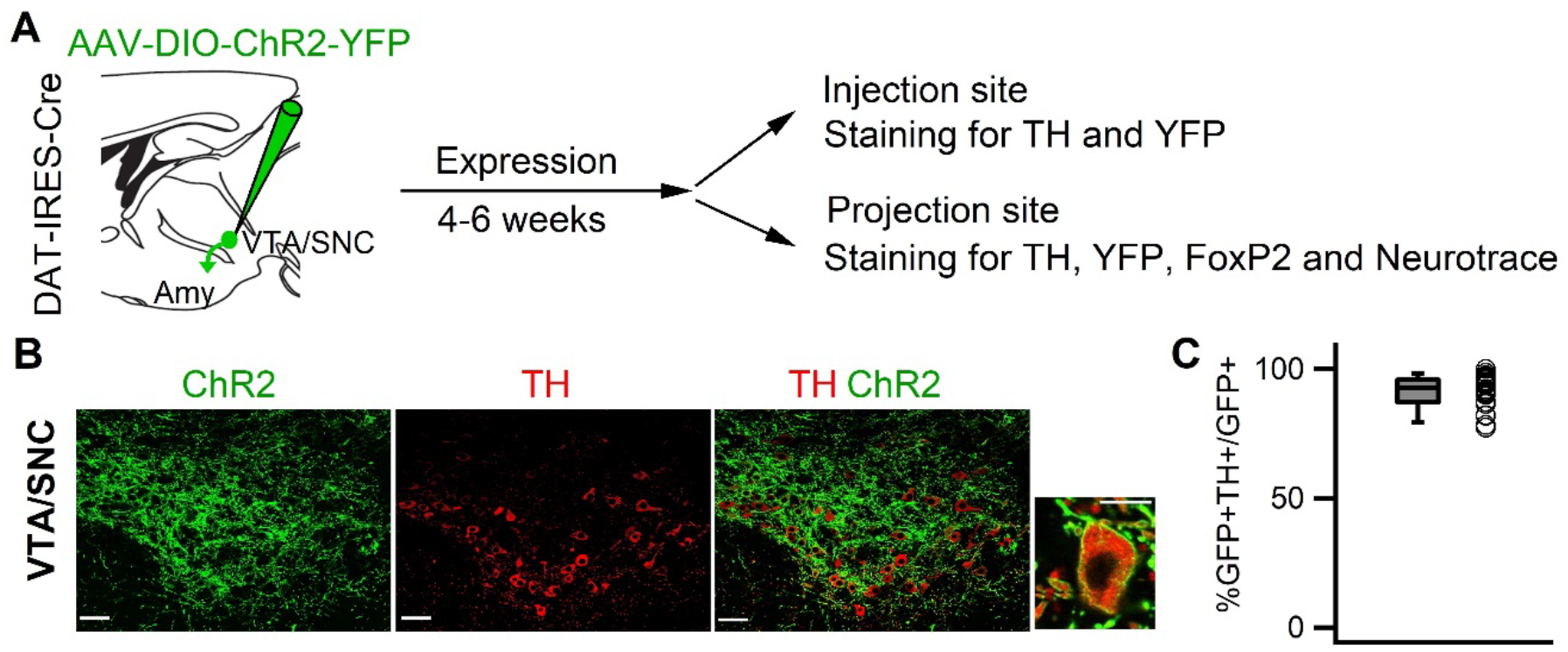
Specific expression of ChR2-YFP in dopaminergic midbrain neurons. **(A)** Experimental strategy: Targeting of dopaminergic midbrain neurons by injecting DAT-IRES-Cre mice with AAV-DIO-ChR2-YFP. Subsequent examination of injection and projection sites using Tyrosine-hydroxylase (TH) and FoxP2 immunostaining, as well as Neurotrace (NT) to delineate brain structures. **(B)** Single plane confocal images (0.9 μm thickness) of VTA/SNC midbrain neurons expressing ChR2-YFP (green, left) immunolabeled for TH (red, middle), and overlay (right), including close up of a TH-positive ChR2-YFP expressing cell. Scale bars: 100 μm (overview), 10 μm (close-up). **(C)** On average, 91.17 ± 1.37% of ChR2-YPF expressing midbrain neurons were TH-positive (n=10 hemispheres from 5 animals).

**Figure 1–figure supplement 2.**
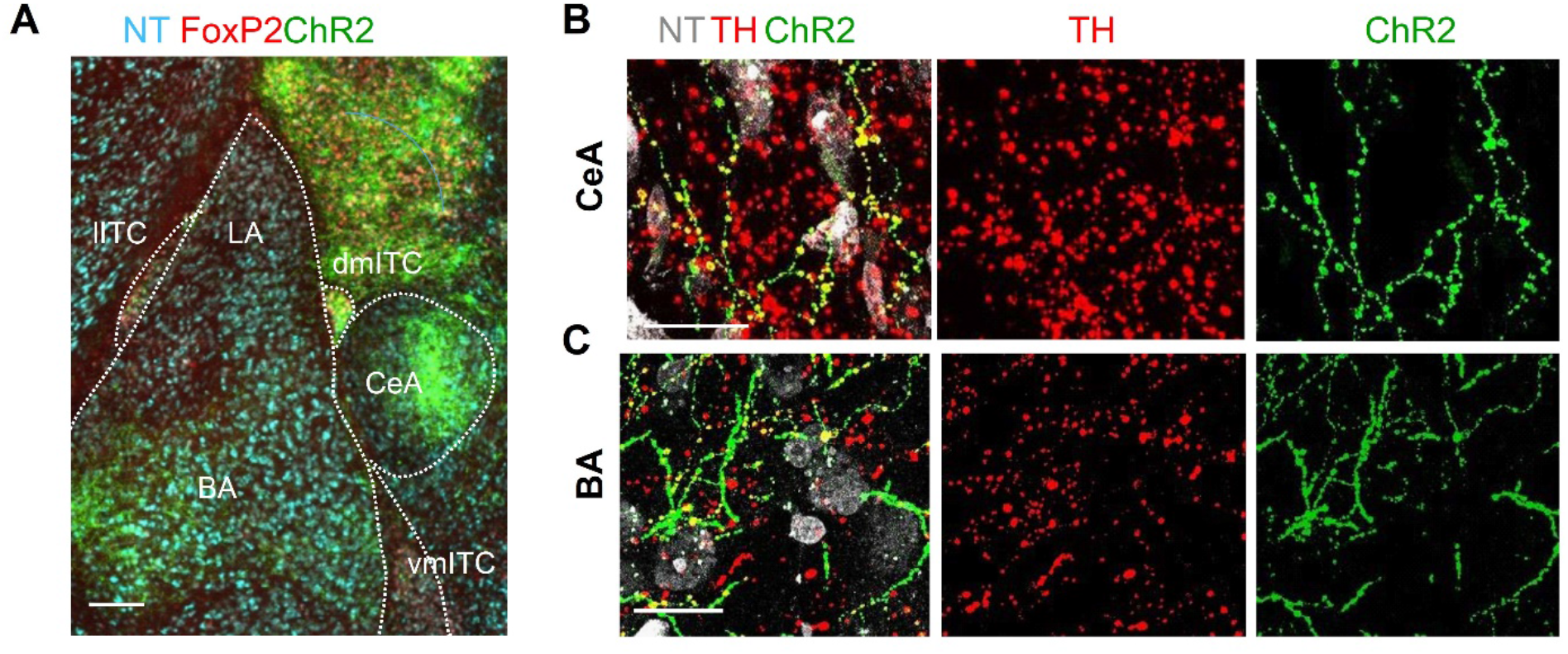
Labeling of TH-positive axons in the amygdala. (A) Fluorescence microscopic overview image of the amygdala showing ChR2-YFP-positive fibers distributed in CeA, BA, ITC clusters, and the amygdala-striatal transition zone. Scale bar: 200 μm. **(B-C)** Maximum intensity projection confocal images of ChR2-YFP-positive fibers (green, right panel), punctate labeling with TH (red) in the same area (middle panel), and overlay together with NT (grey, left panel), demonstrating that ChR2-YFP-labelled fibers are decorated with TH-positive puncta (yellow) in CeA (B) and BLA (C). Thickness of confocal z-stacks: 12.9 and 11.7 μm for and (C), respectively. Scale bars: 10 μm.

**Figure 2–figure supplement 1.**
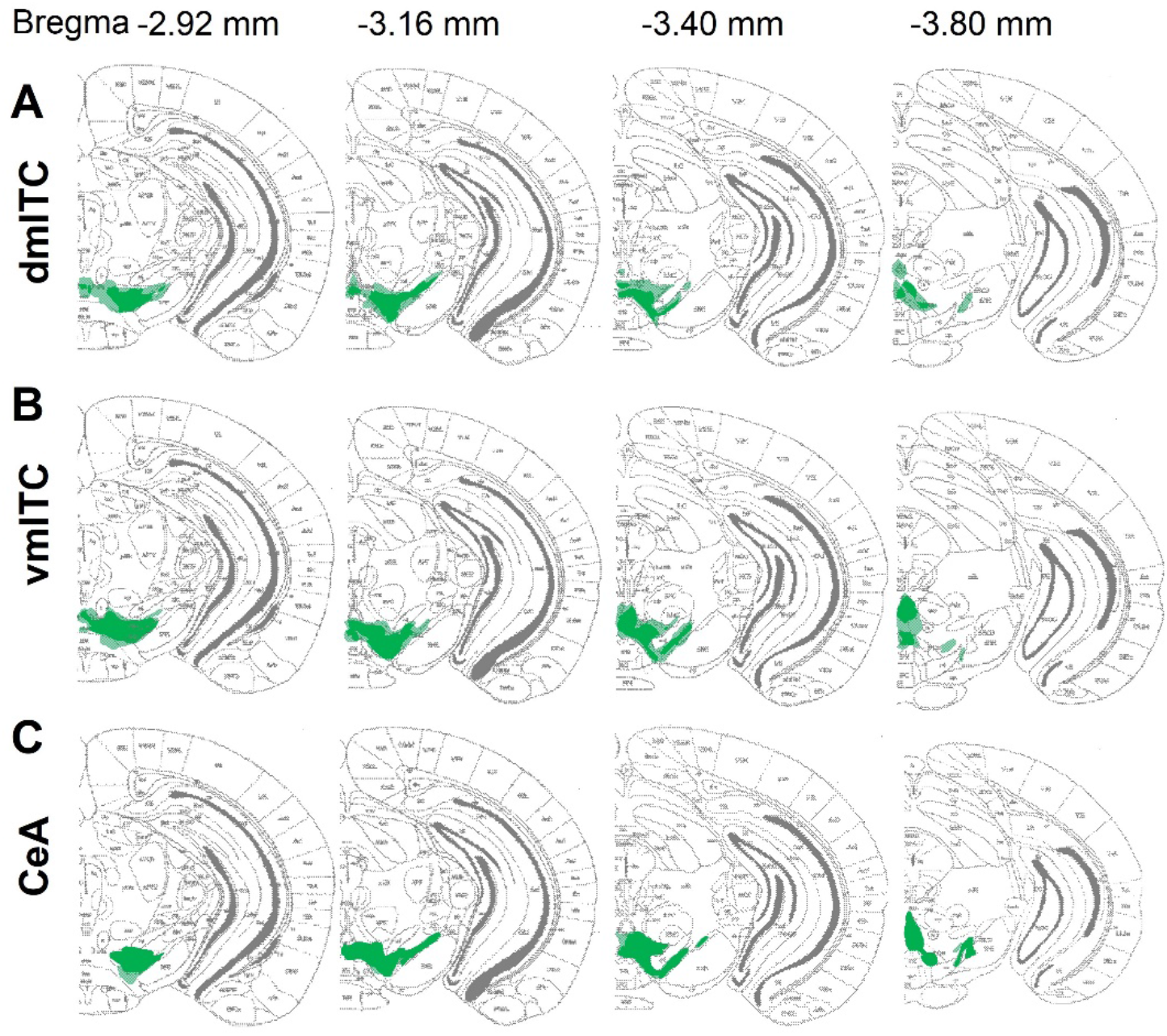
Overview of ChR2-YFP injection and expression sites in the midbrain. Brain atlas overlays of sites showing distribution of ChR2-YFP labelled neuronal cell bodies in the dopaminergic midbrain for all animals in which fast PSCs were recorded. **(A)** Injections sites for dm-ITC recordings (n=16 animals), **(B)** Injections sites for vm-ITC recordings (n=11 animals) and **(C)** Injections sites for CeA recordings (n=11 animals). Bregma levels are indicated at the top.

**Figure 2–figure supplement 2.**
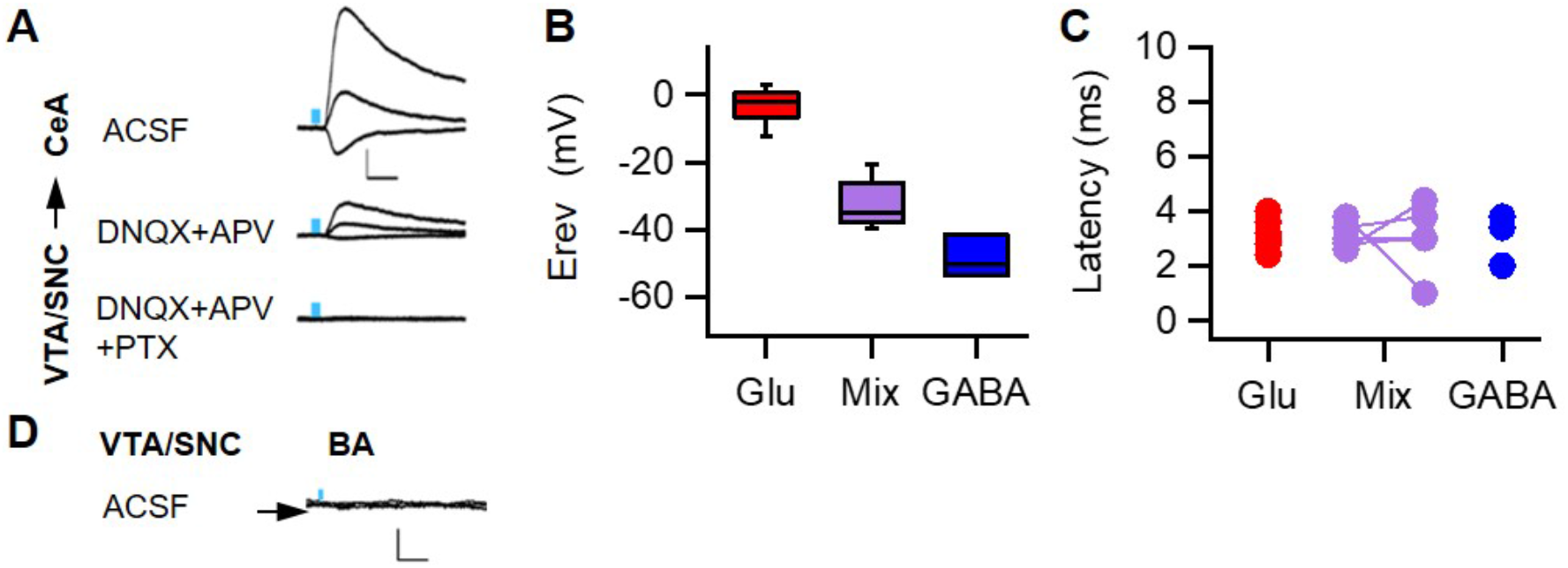
Fast PSCs onto CeA neurons are mainly glutamatergic. **(A)** Example traces of mixed PSCs evoked by dopaminergic fiber stimulation recorded from a CeA neuron at different holding potentials (−70mV, 0mV and 40mV). PSCs were largely blocked with glutamate receptor blockers (20 μM DNQX and 100 μM APV, middle traces) and completely abolished upon addition of the GABA_A_ channel blocker PTX (100 μM, bottom traces). **(B)** Box plot of PSC reversal potentials (Erev) for CeA neurons. Erev for different response types were: Glutamatergic PSCs −2.98 ± 1.54 mV, mixed PSCs −32.75 ± 3.28 mV, and GABAergic −48.58 ± 3.62 mV (one-way ANOVA, F (2, 15) = 90.495, p<0.001). **(C)** Similar latencies of evoked excitatory (glutamatergic 3.14 ± 0.19 ms, n=10), inhibitory (GABAergic 3.07 ± 0.55 ms, n=3) and mixed (glutamatergic component 3.12 ± 0.22 ms and GABA component 3.04 ± 0.57 ms, n=5) PSCs. Scale bars: 40 pA, 10 ms. **(D)** Example traces recorded from a BA projection neuron at different holding potentials (−70 mV, 0 mV and 40 mV) upon dopaminergic fiber stimulation yielding no response (n=10 neurons recorded from 5 animals showing co-release in either CeA or ITCs). Scale bars: 20 pA, 10 ms.

**Figure 2–figure supplement 3.**
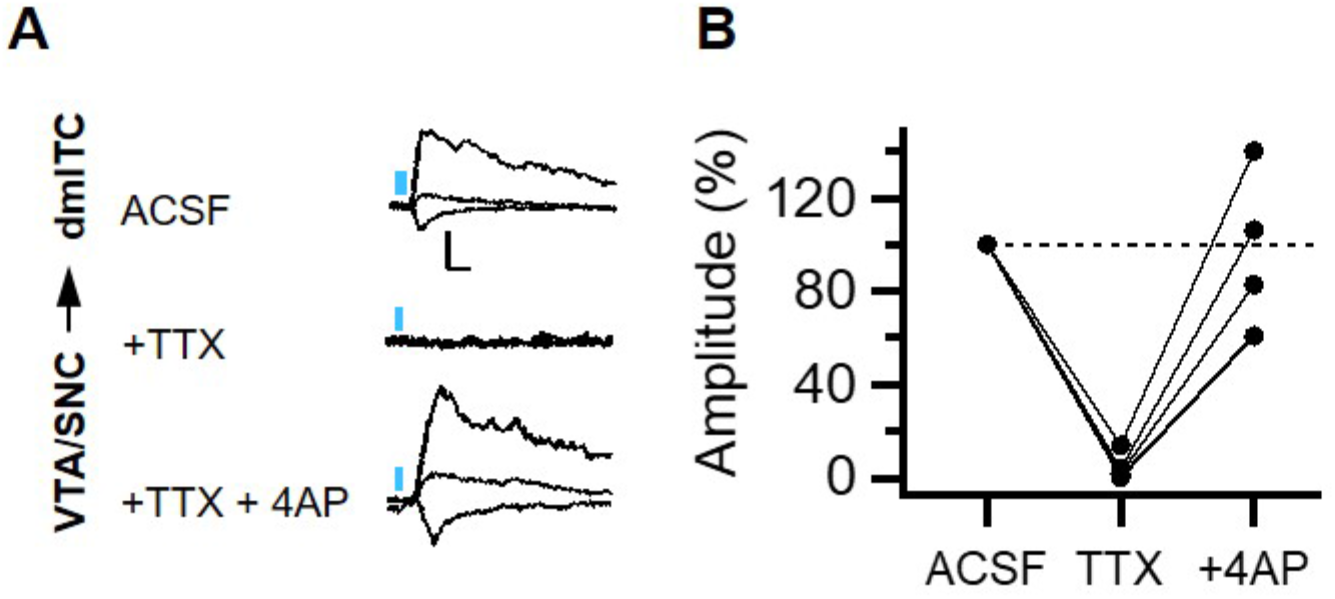
Fast PSCs from midbrain dopaminergic fibers onto dm-ITCs are monosynaptic. **(A)** Example traces of PSCs evoked by dopaminergic fiber stimulation recorded from a dm-ITC at different holding potentials (−70 mV, 0 mV and 40 mV). The PSC is blocked by TTX (0.5 μM) and recovered with 4-AP (100 μM) in the presence of TTX, indicating its monosynaptic nature. Scale bars: 20 pA, 5 ms. **(B)** Summary data of normalized PSC amplitudes for the experiment shown in (A). Repeated measures ANOVA, F (2, 6) =14.88, p=0.013 reveals significant differences between drugs (n=4 dm-ITCs from 3 animals).

**Figure 3–figure supplement 1.**
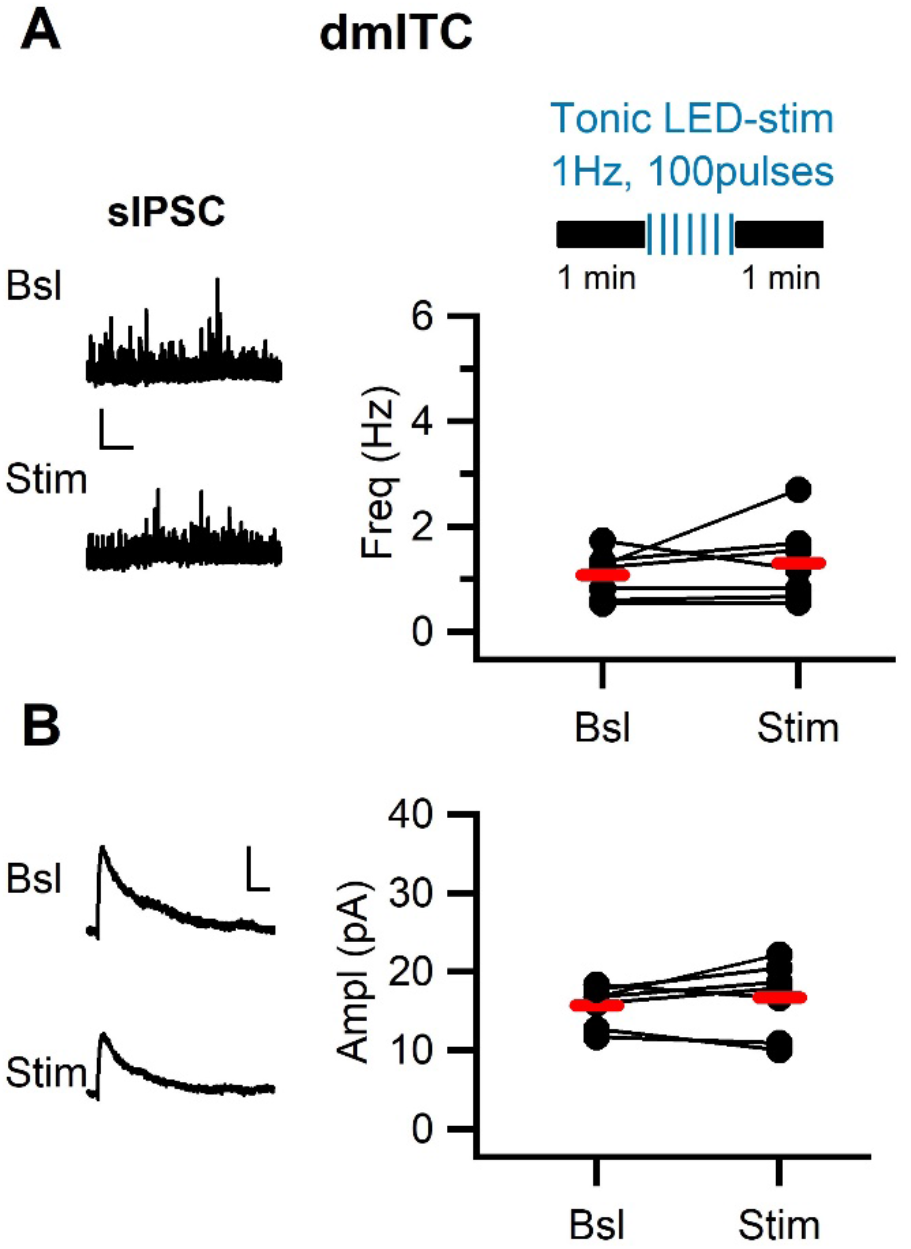
Tonic stimulation of midbrain dopaminergic fibers does not alter sIPSC amplitude in dm-ITCs. **(A)** Top: Experimental protocol. sIPSC were assessed in dmITCs before and after tonic stimulation of dopaminergic fibers (1 Hz,100 pulses). Left: Example traces of sIPSC activity recorded at 0 mV from a dm-ITC before (Bsl) and after stimulation (Stim). Scale bars: 20 pA, 10 s. Right: Plot of sIPSC frequency in individual neurons (dots) and average (red bars), which was not affected by dopaminergic fiber stimulation (1.07 ± 0.17 Hz vs. 1.31 ± 0.28 Hz, n=7, paired t-test p= 0.348). **(B)** Left: Example traces of averaged sIPSC before and after stimulation. Scale bars: 5 pA, 20 ms. Right: Plot of sIPSC amplitude in individual neurons (dots) and average (red bars), which remained stable after stimulation (15.71 ± 0.95 pA vs. 16.68 ± 1.73 pA, n=7, paired t-test p=0.398).

**Figure 3–figure supplement 2.**
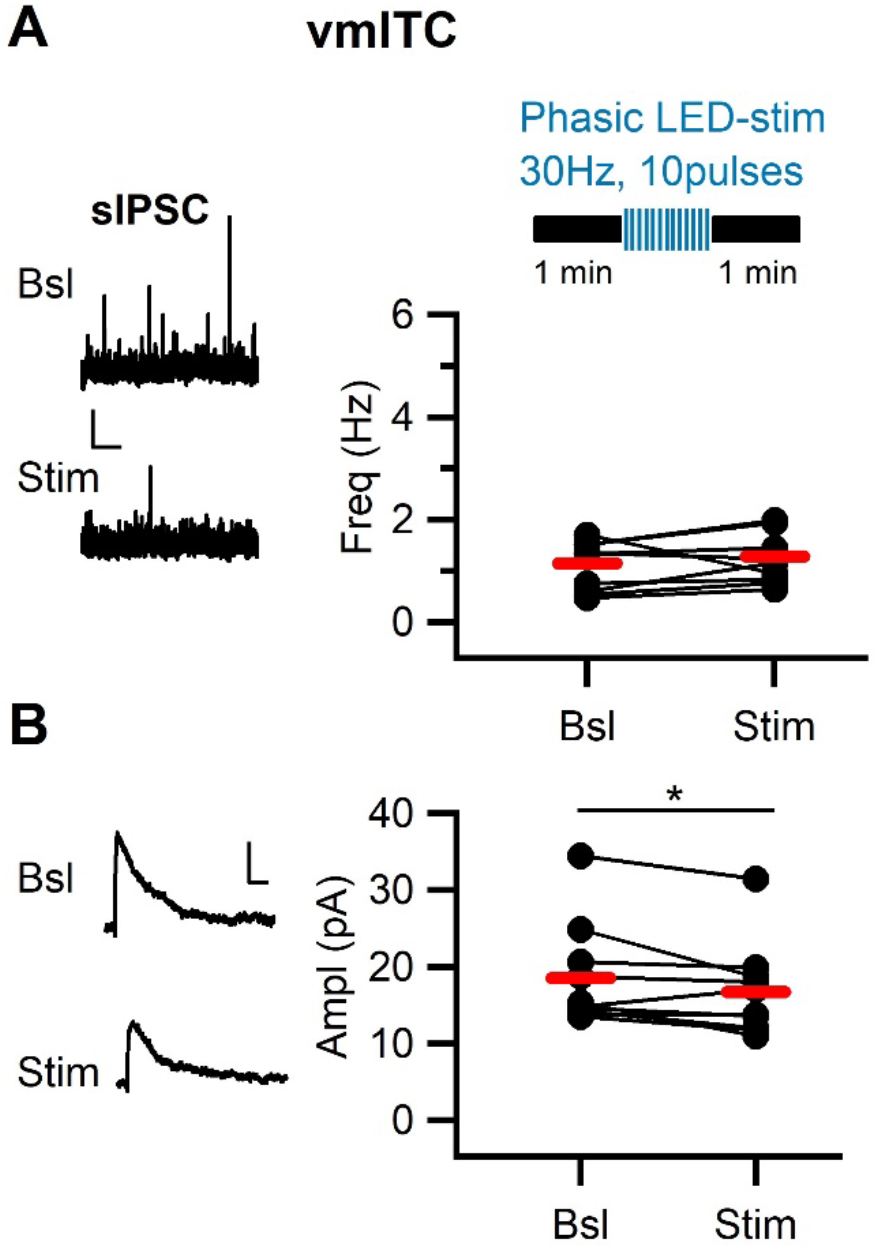
Phasic stimulation of midbrain dopaminergic fibers alters sIPSC amplitude in vm-ITCs. **(A)** Top: Experimental protocol. sIPSCs were assessed in vmITCs before and after phasic stimulation of dopaminergic fibers (30 Hz, 10 pulses, 10 sweeps). Left: Example traces of sIPSC activitiy recorded at 0 mV (A) recorded from vm-ITCs before (Bsl) and after stimulation (Stim). Scale bars: 20 pA, 10s. Right: Plot of sIPSC frequency in individual neurons (dots) and average (red lines). sIPSC frequency (1.14 ± 0.16 Hz vs. 1.28 ± 0.18 Hz, n=10, paired t-test p=0.272) was not affected by dopaminergic fiber stimulation. **(B)** Left: Example traces of averaged sIPSCs before and after stimulation. Scale bars: for sIPSC 5 pA, 20 ms. Right: Plot of sIPSC amplitudes in individual neurons (dots) and average (red lines). sIPSC amplitude decreased after stimulation (18.50 ± 2.12 pA vs. 16.68 ± 2.15 pA, n=10, paired t-test *p=0.034).

**Figure 5–figure supplement 1.**
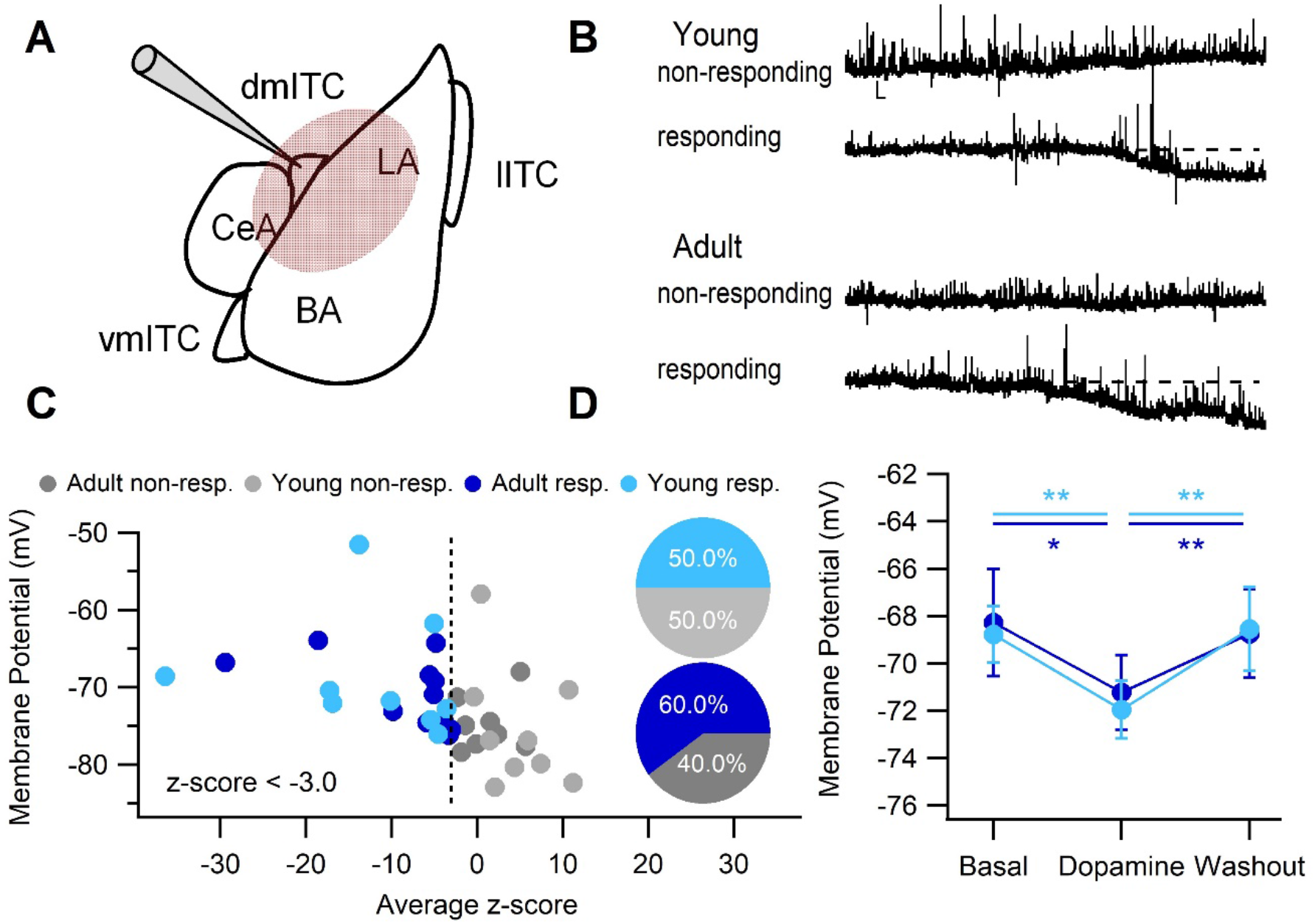
Bath application of DA hyperpolarizes a fraction of dm-ITCs in both young and adult mice. **(A)** Experimental approach: Recording of dm-ITCs and bath application of DA. **(B)** Representative traces of dm-ITC neurons recorded at resting membrane potential in current clamp mode from young (postnatal day 20-28) and adult (8-10 weeks) mice. DA (15 to 30 μM) was applied during the time indicated. Scale bars: 2 mV, 10 s. DA hyperpolarized some cells (responding), but not others (non-responding). **(C)** Distribution of recorded cells according to their response to DA and effect on membrane potential represented as z-scores. Responding cells (z-score cut-off at −3) are shown in blue, non-responding cells in grey. Insets: Fraction of DA-responsive neurons in young mice (top, n=18) and adult mice (bottom, n=20). **(D)** Absolute changes in membrane potential in DA-responsive neurons recorded from young (light blue) and adult (dark blue) mice. DA significantly hyperpolarized responsive dm-ITCs (repeated measures ANOVA for young (n=9): F (2, 16) = 16.23, p<0.001, Bonferroni post-hoc comparison Bsl vs. DA, **p=0.001, DA vs. Wash **p=0.004; for adult (n=11): F (2, 20) = 9.17; p=0.001, Bonferroni post-hoc comparison Bsl vs. DA, *p=0.016, DA vs. Wash **p=0.004).

**Figure 6–figure supplement 1.**
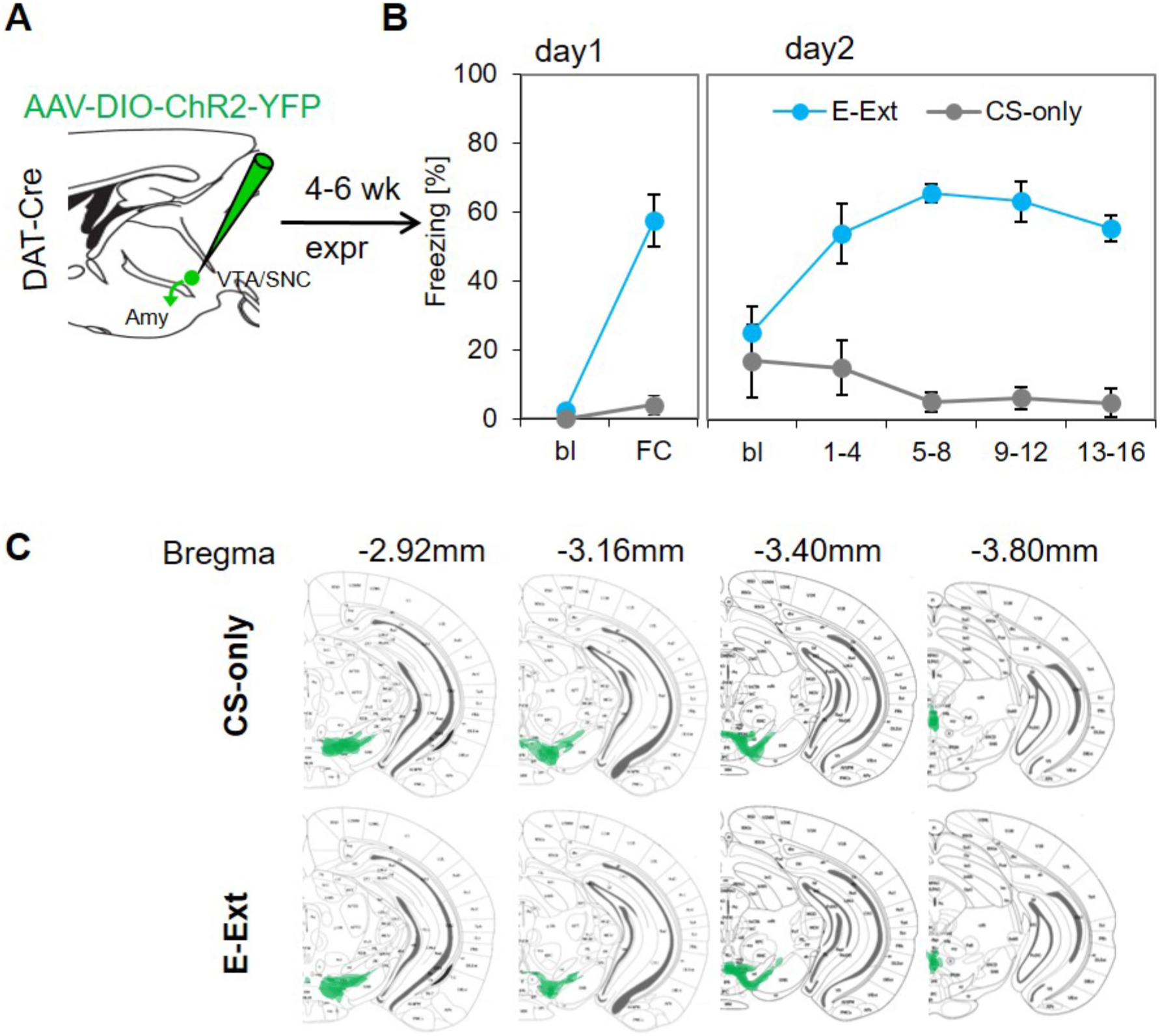
Behavioral data from DAT-IRES-Cre mice used for ex vivo recordings and viral injection sites. **(A)** Experimental strategy for targeting of dopaminergic midbrain neurons by injecting DAT-IRES-Cre mice with AAV-DIO-ChR2-YFP. **(B)** Freezing levels for behavioral paradigms shown in Figure 6A in virally transduced DAT-IRES-Cre mice used for recordings. A group of mice was subjected to only CS presentations (CS-Only, grey, n=4 mice), the other group was subjected to fear conditioning on day 1 and early extinction training with 16 CS presentations (E-Ext, blue, n=6 mice). Repeated measures ANOVA, F (3, 15) = 1.382, p=0.287 reveals no significant differences between time points for E-Ext. **(C)** Brain atlas overlays of sites showing the distribution of ChR2-YFP labelled neuronal cell bodies in the dopaminergic midbrain of DAT-IRES-Cre mice in which fast PSCs were recorded. Top: CS-only (n=4 animals), Bottom: E-Ext (n=6 animals). Bregma levels are indicated at the top.

**Figure 6–figure supplement 2.**
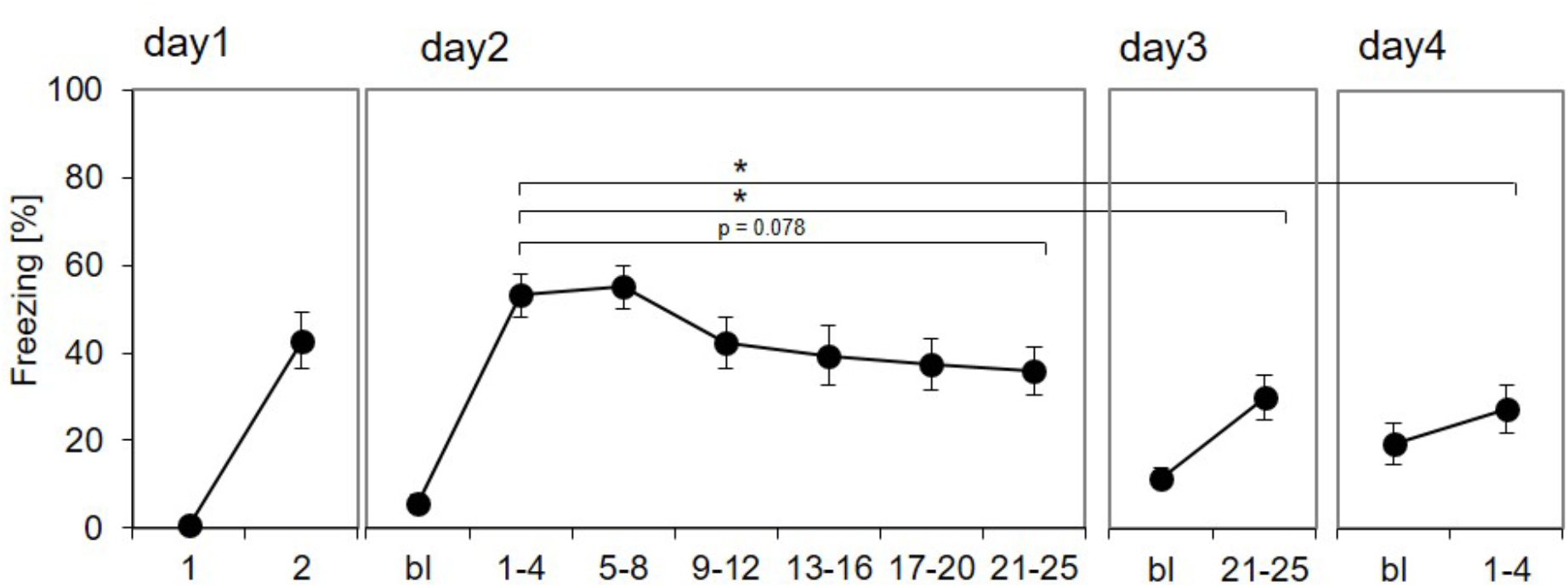
Behavioral data from mice trained with a long extinction protocol. Another group of DAT-IRES-Cre mice was subjected to extinction training with 25 CS presentations on days 2 and 3, and extinction recall by 4 CS presentations on day 4 (n=10 mice). On days 2 and 3, freezing was averaged across 4 CSs for the first 5 time points and 5 CSs for the last one. Repeated measures ANOVA, F (3, 27) = 6.467, p=0.002 reveals significant differences between time points; Bonferroni post-hoc comparison: day1 (CS1-4) vs. day3 (CS21-25) *p=0.031, day1 (CS1-4) vs. day4 (CS1-4) *p=0.037.

**Figure 6–figure supplement 3.**
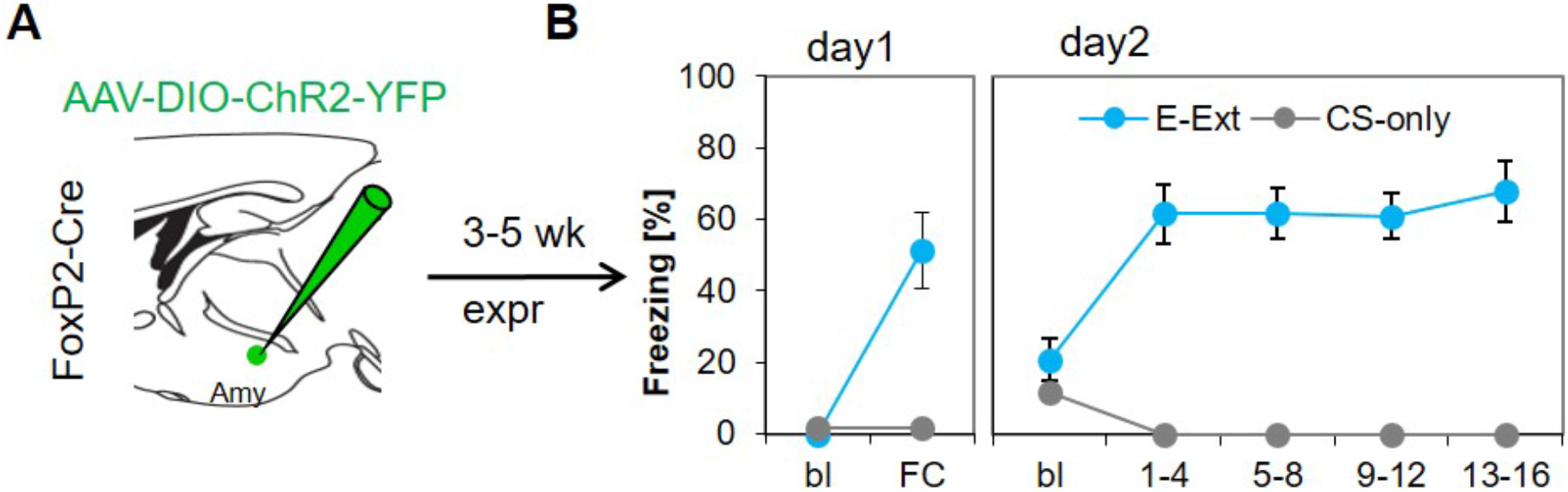
Behavioral data from FoxP2-Cre mice used for ex vivo recordings. **(A)** Experimental strategy for targeting of dm-ITC clusters by injecting FoxP2-Cre mice with AAV-DIO-ChR2-YFP. **(B)** Freezing levels of virally transduced FoxP2-Cre mice used for recordings (CS-Only group, grey, n=4; E-Ext group, blue, n=6) are displayed as described in (Figure 6–figure supplement 1). Repeated measures ANOVA, F (3,15) = 0.647, p=0.597 reveals no significant differences between time points for E-Ext.

## References

Abraham, A. D., Neve, K. A. & Lattal, K. M. 2014. Dopamine and extinction: a convergence of theory with fear and reward circuitry. Neurobiol Learn Mem, 108, 65–77. 10.1016/j.nlm.2013.11.007

Amano, T., Unal, C. T. & Pare, D. 2010. Synaptic correlates of fear extinction in the amygdala. Nat Neurosci, 13, 489–94. 10.1038/nn.2499

Amir, A., Amano, T. & Pare, D. 2011. Physiological identification and infralimbic responsiveness of rat intercalated amygdala neurons. J Neurophysiol, 105, 3054–66. 10.1152/jn.00136.2011

Asan, E. 1997. Ultrastructural features of tyrosine-hydroxylase-immunoreactive afferents and their targets in the rat amygdala. Cell Tissue Res, 288, 449–69. 10.1007/s004410050832

Asede, D., Bosch, D., Luthi, A., Ferraguti, F. & Ehrlich, I. 2015. Sensory inputs to intercalated cells provide fear-learning modulated inhibition to the basolateral amygdala. Neuron, 86, 541–54. 10.1016/j.neuron.2015.03.008

Backman, C. M., Malik, N., Zhang, Y., Shan, L., Grinberg, A., Hoffer, B. J., Westphal, H. & Tomac, A. C. 2006. Characterization of a mouse strain expressing Cre recombinase from the 3’ untranslated region of the dopamine transporter locus. Genesis, 44, 383–90. 10.1002/dvg.20228

Barsy, B., Kocsis, K., Magyar, A., Babiczky, A., Szabo, M., Veres, J. M., Hillier, D., Ulbert, I., Yizhar, O. & Matyas, F. 2020. Associative and plastic thalamic signaling to the lateral amygdala controls fear behavior. Nat Neurosci, 23, 625–637. 10.1038/s41593-020-0620-z

Belujon, P. & Grace, A. A. 2015. Regulation of dopamine system responsivity and its adaptive and pathological response to stress. Proc Biol Sci, 282. 10.1098/rspb.2014.2516

Bouton, M. E. 2004. Context and behavioral processes in extinction. Learn Mem, 11, 485–94. 10.1101/lm.78804

Brischoux, F., Chakraborty, S., Brierley, D. I. & Ungless, M. A. 2009. Phasic excitation of dopamine neurons in ventral VTA by noxious stimuli. Proc Natl Acad Sci U S A, 106, 4894–9. 10.1073/pnas.0811507106

Bromberg-Martin, E. S., Matsumoto, M. & Hikosaka, O. 2010. Dopamine in motivational control: rewarding, aversive, and alerting. Neuron, 68, 815–34. 10.1016/j.neuron.2010.11.022

Busti, D., Geracitano, R., Whittle, N., Dalezios, Y., Manko, M., Kaufmann, W., Satzler, K., Singewald, N., Capogna, M. & Ferraguti, F. 2011. Different fear states engage distinct networks within the intercalated cell clusters of the amygdala. J Neurosci, 31, 5131–5144. 10.1523/JNEUROSCI.6100-10.2011

Cho, J. H., Deisseroth, K. & Bolshakov, V. Y. 2013. Synaptic encoding of fear extinction in mPFC-amygdala circuits. Neuron, 80, 1491–1507. 10.1016/j.neuron.2013.09.025

Davis, M. 2000. The role of the amygdala in conditioned and unconditioned fear and anxiety. *In:* Aggleton, J. P. (ed.) The Amygdala. Oxford University Press.

Duvarci, S. & Pare, D. 2014. Amygdala microcircuits controlling learned fear. Neuron, 82, 966–80. 10.1016/j.neuron.2014.04.042

Ehrlich, I., Humeau, Y., Grenier, F., Ciocchi, S., Herry, C. & Luthi, A. 2009. Amygdala inhibitory circuits and the control of fear memory. Neuron, 62, 757–771. 10.1016/j.neuron.2009.05.026

Ferrazzo, S., Gunduz-Cinar, O., Stefanova, N., Pollack, G. A., Holmes, A., Schmuckermair, C. & Ferraguti, F. 2019. Increased anxiety-like behavior following circuit-specific catecholamine denervation in mice. Neurobiol Dis, 125, 55–66. 10.1016/j.nbd.2019.01.009

Fuxe, K., Jacobsen, K. X., Hoistad, M., Tinner, B., Jansson, A., Staines, W. A. & Agnati, L. F. 2003. The dopamine D1 receptor-rich main and paracapsular intercalated nerve cell groups of the rat amygdala: relationship to the dopamine innervation. Neuroscience, 119, 733–46. 10.1016/s0306-4522(03)00148-9

Geracitano, R., Kaufmann, W. A., Szabo, G., Ferraguti, F. & Capogna, M. 2007. Synaptic heterogeneity between mouse paracapsular intercalated neurons of the amygdala. J Physiol, 585, 117–34. 10.1113/jphysiol.2007.142570

Grace, A. A., Floresco, S. B., Goto, Y. & Lodge, D. J. 2007. Regulation of firing of dopaminergic neurons and control of goal-directed behaviors. Trends Neurosci, 30, 220–7. 10.1016/j.tins.2007.03.003

Granger, A. J., Wallace, M. L. & Sabatini, B. L. 2017. Multi-transmitter neurons in the mammalian central nervous system. Curr Opin Neurobiol, 45, 85–91. 10.1016/j.conb.2017.04.007

Gregoriou, G. C., Kissiwaa, S. A., Patel, S. D. & Bagley, E. E. 2019. Dopamine and opioids inhibit synaptic outputs of the main island of the intercalated neurons of the amygdala. Eur J Neurosci, 50, 2065–2074. 10.1111/ejn.14107

Groessl, F., Munsch, T., Meis, S., Griessner, J., Kaczanowska, J., Pliota, P., Kargl, D., Badurek, S., Kraitsy, K., Rassoulpour, A., Zuber, J., Lessmann, V. & Haubensak, W. 2018. Dorsal tegmental dopamine neurons gate associative learning of fear. Nat Neurosci, 21, 952–962. 10.1038/s41593-018-0174-5

Guarraci, F. A., Frohardt, R. J. & Kapp, B. S. 1999. Amygdaloid D1 dopamine receptor involvement in Pavlovian fear conditioning. Brain Res, 827, 28–40. S0006-8993(99)01291-3 [pii]

Herry, C. & Johansen, J. P. 2014. Encoding of fear learning and memory in distributed neuronal circuits. Nat Neurosci, 17, 1644–54. 10.1038/nn.3869

Hikind, N. & Maroun, M. 2008. Microinfusion of the D1 receptor antagonist, SCH23390 into the IL but not the BLA impairs consolidation of extinction of auditory fear conditioning. Neurobiol. Learn. Mem., 90. 10.1016/j.nlm.2008.03.003

Holloway, Z. R., Freels, T. G., Comstock, J. F., Nolen, H. G., Sable, H. J. & Lester, D. B. 2018. Comparing phasic dopamine dynamics in the striatum, nucleus accumbens, amygdala, and medial prefrontal cortex. Synapse, e22074. 10.1002/syn.22074

Horvitz, J. C. 2000. Mesolimbocortical and nigrostriatal dopamine responses to salient non-reward events. Neuroscience, 96, 651–6.

Huang, C. C., Chen, C. C., Liang, Y. C. & Hsu, K. S. 2014. Long-term potentiation at excitatory synaptic inputs to the intercalated cell masses of the amygdala. Int J Neuropsychopharmacol, 17, 1233–42. 10.1017/S1461145714000133

Inglis, F. M. & Moghaddam, B. 1999. Dopaminergic innervation of the amygdala is highly responsive to stress. J Neurochem, 72, 1088–94. 10.1046/j.1471-4159.1999.0721088.x

Kim, J. I., Ganesan, S., Luo, S. X., Wu, Y. W., Park, E., Huang, E. J., Chen, L. & Ding, J. B. 2015. Aldehyde dehydrogenase 1a1 mediates a GABA synthesis pathway in midbrain dopaminergic neurons. Science, 350, 102–6. 10.1126/science.aac4690

Kuerbitz, J., Arnett, M., Ehrman, S., Williams, M. T., Vorhees, C. V., Fisher, S. E., Garratt, A. N., Muglia, L. J., Waclaw, R. R. & Campbell, K. 2018. Loss of Intercalated Cells (ITCs) in the Mouse Amygdala of Tshz1 Mutants Correlates with Fear, Depression, and Social Interaction Phenotypes. J Neurosci, 38, 1160–1177. 10.1523/JNEUROSCI.1412-17.2017

Kwon, O. B., Lee, J. H., Kim, H. J., Lee, S., Lee, S., Jeong, M. J., Kim, S. J., Jo, H. J., Ko, B., Chang, S., Park, S. K., Choi, Y. B., Bailey, C. H., Kandel, E. R. & Kim, J. H. 2015. Dopamine Regulation of Amygdala Inhibitory Circuits for Expression of Learned Fear. Neuron, 88, 378–89. 10.1016/j.neuron.2015.09.001

Lamont, E. W. & Kokkinidis, L. 1998. Infusion of the dopamine D1 receptor antagonist SCH 23390 into the amygdala blocks fear expression in a potentiated startle paradigm. Brain Res, 795, 128–36. 10.1016/s0006-8993(98)00281-9

Lee, J. H., Lee, S. & Kim, J. H. 2017. Amygdala Circuits for Fear Memory: A Key Role for Dopamine Regulation. Neuroscientist, 23, 542–553. 10.1177/1073858416679936

Likhtik, E., Popa, D., Apergis-Schoute, J., Fidacaro, G. A. & Pare, D. 2008. Amygdala intercalated neurons are required for expression of fear extinction. Nature, 454, 642–5. 10.1038/nature07167

Liu, C., Kershberg, L., Wang, J., Schneeberger, S. & Kaeser, P. S. 2018. Dopamine Secretion Is Mediated by Sparse Active Zone-like Release Sites. Cell, 172, 706–718 e15. 10.1016/j.cell.2018.01.008

Luo, R., Uematsu, A., Weitemier, A., Aquili, L., Koivumaa, J., McHugh, T. J. & Johansen, J. P. 2018. A dopaminergic switch for fear to safety transitions. Nat Commun, 9, 2483. 10.1038/s41467-018-04784-7

Lutas, A., Kucukdereli, H., Alturkistani, O., Carty, C., Sugden, A. U., Fernando, K., Diaz, V., Flores-Maldonado, V. & Andermann, M. L. 2019. State-specific gating of salient cues by midbrain dopaminergic input to basal amygdala. Nat Neurosci, 22, 1820–1833. 10.1038/s41593-019-0506-0

Manko, M., Geracitano, R. & Capogna, M. 2011. Functional connectivity of the main intercalated nucleus of the mouse amygdala. J Physiol, 589, 1911–25. 10.1113/jphysiol.2010.201475

Marowsky, A., Yanagawa, Y., Obata, K. & Vogt, K. E. 2005. A specialized subclass of interneurons mediates dopaminergic facilitation of amygdala function. Neuron, 48, 1025–37. 10.1016/j.neuron.2005.10.029

McNally, G. P., Johansen, J. P. & Blair, H. T. 2011. Placing prediction into the fear circuit. Trends Neurosci, 34, 283–92. 10.1016/j.tins.2011.03.005

Mingote, S., Chuhma, N., Kusnoor, S. V., Field, B., Deutch, A. Y. & Rayport, S. 2015. Functional Connectome Analysis of Dopamine Neuron Glutamatergic Connections in Forebrain Regions. J Neurosci, 35, 16259–71. 10.1523/JNEUROSCI.1674-15.2015

Morozov, A., Sukato, D. & Ito, W. 2011. Selective suppression of plasticity in amygdala inputs from temporal association cortex by the external capsule. J Neurosci, 31, 339–345. 10.1523/JNEUROSCI.5537-10.2011

Nader, K. & LeDoux, J. E. 1999. Inhibition of the mesoamygdala dopaminergic pathway impairs the retrieval of conditioned fear associations. Behav Neurosci, 113, 891–901. 10.1037//0735-7044.113.5.891

Nicola, S. M. & Malenka, R. C. 1997. Dopamine depresses excitatory and inhibitory synaptic transmission by distinct mechanisms in the nucleus accumbens. J Neurosci, 17, 5697–710.

Pare, D., Quirk, G. J. & Ledoux, J. E. 2004. New vistas on amygdala networks in conditioned fear. J Neurophysiol, 92, 1–9. 10.1152/jn.00153.2004

Pare, D., Royer, S., Smith, Y. & Lang, E. J. 2003. Contextual inhibitory gating of impulse traffic in the intra-amygdaloid network. Ann N Y Acad Sci, 985, 78–91. 10.1111/j.1749-6632.2003.tb07073.x

Pearce, J. M. & Hall, G. 1980. A model for Pavlovian learning: variations in the effectiveness of conditioned but not of unconditioned stimuli. Psych Rev, 87, 532–552.

Pennartz, C. M., Dolleman-Van der Weel, M. J., Kitai, S. T. & Lopes da Silva, F. H. 1992. Presynaptic dopamine D1 receptors attenuate excitatory and inhibitory limbic inputs to the shell region of the rat nucleus accumbens studied in vitro. J Neurophysiol, 67, 1325–34. 10.1152/jn.1992.67.5.1325

Petreanu, L., Mao, T., Sternson, S. M. & Svoboda, K. 2009. The subcellular organization of neocortical excitatory connections. Nature, 457, 1142–1145. 10.1038/nature07709

Pinto, A. & Sesack, S. R. 2008. Ultrastructural analysis of prefrontal cortical inputs to the rat amygdala: spatial relationships to presumed dopamine axons and D1 and D2 receptors. Brain structure & function, 213, 159–75. 10.1007/s00429-008-0180-6

Rescorla, R. A. & Wagner, A. R. (eds.) 1972. A theory of Pavlovian conditioning: variations in the effectiveness of reinforcement and nonreinforcement: Appleton Century Crofts.

Rothman, J. S. & Silver, R. A. 2018. NeuroMatic: An Integrated Open-Source Software Toolkit for Acquisition, Analysis and Simulation of Electrophysiological Data. Front Neuroinform, 12, 14. 10.3389/fninf.2018.00014

Rousso, D. L., Qiao, M., Kagan, R. D., Yamagata, M., Palmiter, R. D. & Sanes, J. R. 2016. Two Pairs of ON and OFF Retinal Ganglion Cells Are Defined by Intersectional Patterns of Transcription Factor Expression. Cell Rep, 15, 1930–44. 10.1016/j.celrep.2016.04.069

Royer, S., Martina, M. & Pare, D. 2000. Polarized synaptic interactions between intercalated neurons of the amygdala. J Neurophysiol, 83, 3509–18. 10.1152/jn.2000.83.6.3509

Salinas-Hernandez, X. I., Vogel, P., Betz, S., Kalisch, R., Sigurdsson, T. & Duvarci, S. 2018. Dopamine neurons drive fear extinction learning by signaling the omission of expected aversive outcomes. Elife, 7. 10.7554/eLife.38818

Schultz, W., Dayan, P. & Montague, P. R. 1997. A neural substrate of prediction and reward. Science, 275, 1593–9. 10.1126/science.275.5306.1593

Strobel, C., Marek, R., Gooch, H. M., Sullivan, R. K. & Sah, P. 2015. Prefrontal and Auditory Input to Intercalated Neurons of the Amygdala. Cell Rep, 10, 1435–1442. 10.1016/j.celrep.2015.02.008

Stuber, G. D., Hnasko, T. S., Britt, J. P., Edwards, R. H. & Bonci, A. 2010. Dopaminergic terminals in the nucleus accumbens but not the dorsal striatum corelease glutamate. J Neurosci, 30, 8229–33. 10.1523/JNEUROSCI.1754-10.2010

Tamamaki, N., Yanagawa, Y., Tomioka, R., Miyazaki, J., Obata, K. & Kaneko, T. 2003. Green fluorescent protein expression and colocalization with calretinin, parvalbumin, and somatostatin in the GAD67-GFP knock-in mouse. J Comp Neurol, 467, 60–79. 10.1002/cne.10905

Taverna, S., Canciani, B. & Pennartz, C. M. 2005. Dopamine D1-receptors modulate lateral inhibition between principal cells of the nucleus accumbens. J Neurophysiol, 93, 1816–9. 10.1152/jn.00672.2004

Tecuapetla, F., Patel, J. C., Xenias, H., English, D., Tadros, I., Shah, F., Berlin, J., Deisseroth, K., Rice, M. E., Tepper, J. M. & Koos, T. 2010. Glutamatergic signaling by mesolimbic dopamine neurons in the nucleus accumbens. J Neurosci, 30, 7105–10. 10.1523/JNEUROSCI.0265-10.2010

Tritsch, N. X., Ding, J. B. & Sabatini, B. L. 2012. Dopaminergic neurons inhibit striatal output through non-canonical release of GABA. Nature, 490, 262–6. 10.1038/nature11466

Tritsch, N. X., Granger, A. J. & Sabatini, B. L. 2016. Mechanisms and functions of GABA co-release. Nat Rev Neurosci, 17, 139–45. 10.1038/nrn.2015.21

Tritsch, N. X., Oh, W. J., Gu, C. & Sabatini, B. L. 2014. Midbrain dopamine neurons sustain inhibitory transmission using plasma membrane uptake of GABA, not synthesis. Elife, 3, e01936. 10.7554/eLife.01936

Tsai, H. C., Zhang, F., Adamantidis, A., Stuber, G. D., Bonci, A., de Lecea, L. & Deisseroth, K. 2009. Phasic firing in dopaminergic neurons is sufficient for behavioral conditioning. Science, 324, 1080–4. 10.1126/science.1168878

Wei, X., Ma, T., Cheng, Y., Huang, C. C. Y., Wang, X., Lu, J. & Wang, J. 2018. Dopamine D1 or D2 receptor-expressing neurons in the central nervous system. Addict Biol, 23, 569–584. 10.1111/adb.12512

Yokoyama, M., Suzuki, E., Sato, T., Maruta, S., Watanabe, S. & Miyaoka, H. 2005. Amygdalic levels of dopamine and serotonin rise upon exposure to conditioned fear stress without elevation of glutamate. Neurosci Lett, 379, 37–41. 10.1016/j.neulet.2004.12.047

